# Molecular interaction mechanism of a 14-3-3 protein with a phosphorylated peptide elucidated by enhanced conformational sampling

**DOI:** 10.1101/2020.05.24.113209

**Authors:** Junichi Higo, Takeshi Kawabata, Ayumi Kusaka, Kota Kasahara, Narutoshi Kamiya, Ikuo Fukuda, Kentaro Mori, Yutaka Hata, Yoshifumi Fukunishi, Haruki Nakamura

## Abstract

Enhanced conformational sampling, a genetic-algorithm-guided multi-dimensional virtual-system coupled molecular dynamics, can provide equilibrated conformational distributions of a receptor protein and a flexible ligand at room temperature. The distributions provide not only the most stable but also semi-stable complex structures, and propose a ligand–receptor binding process. This method was applied to a system consisting of a receptor protein, 14-3-3ε, and a flexible peptide, phosphorylated Myeloid leukemia factor 1 (pMLF1). The results present comprehensive binding pathways of pMLF1 to 14-3-3ε. We identified four thermodynamically stable clusters of MLF1 on the 14-3-3ε surface, and free-energy barriers among some clusters. The most stable cluster includes two high-density spots connected by a narrow corridor. When pMLF1 passes the corridor, a salt-bridge relay (switching) related to the phosphorylated residue of pMLF1 occurs. Conformations in one high-density spots are similar to the experimentally determined complex structure. Three-dimensional distributions of residues in the intermolecular interface rationally explain the binding-constant changes resultant from alanine–mutation experiment for the residues. We performed a simulation of non-phosphorylated peptide and 14-3-3ε, which demonstrated that the complex structure was unstable, suggesting that phosphorylation of the peptide is crucially important for binding to 14-3-3ε.

## INTRODUCTION

The 14-3-3 proteins are eukaryotic adapter proteins that bind short phosphorylated sequences (14-3-3 interaction sequences).^1–3^ They bind to numerous partner proteins, usually by recognition of a phosphoserine or phosphothreonine motif. Although several binding motif patterns have been proposed for 14-3-3 interaction sequences, the mechanism of their binding specificities is not understood clearly yet, except for salt bridges between the phosphorylated site of 14-3-3 interaction sequences and 14-3-3 proteins.

Binding of 14-3-3 proteins and 14-3-3 interaction sequences can regulate the subcellular localization and enzymatic activities of proteins related to signal transduction and cell-cycle regulation.^4^ Therefore, 14-3-3 proteins are well known as potential drug targets. Small molecules such as peptide mimetic and natural products have been explored to elucidate inhibitors and stabilizers of 14-3-3 protein complexes.^5^

Humans have seven 14-3-3 proteins that are mutually similar: *β, γ, ε, η, σ, τ,* and *ζ*. Among them, the sequence identities are higher than 60%. After we analyzed the 14-3-3 interaction sequences that bind to the human 14-3-3ε and *ζ* proteins using MATRAS software,^6^ we presented the aligned sequence in Figures S1A and S1B of Supporting Information (SI). The figures portray S1C and S1D, displaying the superimposed structures of the 14-3-3 interaction sequences binding to 14-3-3ε and 14-3-3 ζ proteins from two orientations. Figure S1E shows the RMSD of the superimposed structures at each residue site. Although some diversity exists in the sequences (Figures S1A and S1B), Figures S1C–E show that the binding structures of the 14-3-3 interaction sequences have similarity around the phosphorylated residue. Therefore, if the complex-formation mechanisms of a 14-3-3 protein and an interaction sequence are understood using a computational method, then this mechanism would be useful for understanding other 14-3-3 complexes.

Molzan et al. solved an X-ray crystal structure (PDB ID: 3ual) (Figure S2) of the human 14-3-3ε protein bound to a phosphorylated peptide (sequence = RSFpSEPFG, where pS is a phosphorylated serine at the 34th residue position) taken from myeloid leukemia factor 1 (MLF1).^7^ They also measured the dissociation constants *K_D_* of alanine-mutated MLF1 peptides at the 33rd, 35th, and 37th residues (F33A, E35A, and F37A) using isothermal titration calorimetry (ITC). It is particularly interesting that only the mutation F33A led to a significant decrease in affinity, which might be attributable to the loss of hydrophobic interactions by Phe 33 sandwiched between two hydrophobic residues Leu 223 and Leu 230 of the 14-3-3ε protein (Figure S2). In contrast, E35A and F37A exhibited almost no change in *K_D_*. Phe 37 is located on the surface of 14-3-3ε in the complex forming a weak hydrophobic contact to the stem of Lys 40 of 14-3-3ε. This can be a reason for the negligibly small change of *K_D_* by F37A. In the crystal structure, however, Glu 35 is located at a position that can form salt bridges to Lys 50 and Lys 123 of 14-3-3ε. Therefore, explaining the reason for the small change of *K_D_* by E35A based on the crystal structure is difficult.

The 14-3-3 proteins are related to many human diseases. Kimura et al. reported that a mutation of nuclear distribution E homolog 1 (NDE1) has significant association with schizophrenia.^8^ The 14-3-3ε protein bound to the mutated site of NDE1. The 14-3-3 proteins bind to myeloid leukemia factor 1 (MLF1), which is reportedly overexpressed in acute myeloid leukemiaΛ^11^ Although Winteringham et al. (2006) reported that the nuclear content of mouse MLF1 was regulated by 14-3-3 binding,^11^ Molzan and Ottmann (2013) reported that the subcellular localization of human MLF1 is independent of 14-3-3 proteins.^12^ Therefore, the biological role of 14-3-3ε/MLF1 remains unknown. However, this complex is an appropriate system for the study of complex–formation processes between 14-3-3 proteins and the 14-3-3 interaction sequences comprehensively by computations because, as described above, the tertiary structure of 14-3-3ε/MLF1 was well studied. The amino-acid contribution to the complex stability was also measured.^7^ We infer that, if the computations can explain the measured amino-acid contribution to the complex stability consistently, then the comprehensive binding mechanism is reliable.

Generalized ensemble methods such as the multicanonical sampling,^13–16^ replica exchange^17,18^ method, and their variants^19–25^ are powerful methods to sample large-scale motions of a molecular system by enhancing the conformational motions along the energy or temperature axis. However, these methods might overlook less-stable conformations (i.e., conformations in minor basins) when the minor basins overlap to the major basins in the axis.^26,27^ This oversight is not problematic when the minor basins are beyond the scope of the study. However, when the minor basins act as bridges to connect the major basins, then the oversight might cause conformation trapping, which hinders sampling of a wide conformational space. Consequently, an alternative computational approach must be used.

Adaptive umbrella sampling^28,29^ might avoid such an oversight or trapping because one can set a reaction coordinate so that the major and minor basins are discriminated along the reaction-coordinate axis. However, this method requires fine tuning of a weight function (bias function). Practically, the difficulty of tuning might require a very long simulation (or increment of the number of iterative simulations) to treat a complicated system.

Recently, we introduced a method, virtual-system coupled sampling method,^30^ and combined it to multicanonical^31^ and adaptive umbrella sampling.^26,32^ These methods provide an equilibrated conformational ensemble of the system. Nevertheless, they still require fine tuning of the bias function. We introduced a new method, multi-dimensional virtual-system coupled molecular dynamics (mD-VcMD),^38,39^ which is free from fine tuning of the bias function. Furthermore, we extended mD-VcMD using a genetic algorithm and designated it as genetic algorithm-guided mD-VcMD (GA-guided mD-VcMD).^40^

For this study, we performed GA-guided mD-VcMD simulations of a system consisting of the receptor (14-3-3ε protein) and the ligand (MLF1 segment) to obtain the free-energy landscape which controls ligand–receptor binding. MLF1 was positioned distant from the binding site of 14-3-3ε in the initial conformations of simulation. MLF1 of two types were examined: phosphorylated and non-phosphorylated MLF1. We examined the contributions of Phe 33, Sep 34, Glu 35, and Phe 37 of MLF1 to the complex stability by analyzing conformational distributions obtained from the simulation, and calculated the free-energy landscape (3D conformational distribution) to discuss comprehensive binding processes of MLF1 to 14-3-3ε.

## METHODS

### Conformational ensemble from GA-guided mD-VcMD

To sample both the unbound and bound states of the receptor-ligand system, we use one of generalized ensemble methods: GA-guided mD-VcMD, which is an extension of mD-VcMD^38,39^. Methodological details are explained elsewhere,^40^ and actual parameter values used for the simulation are described in section 4 of SI. Here, only a brief outline of the method is introduced below. This method controls the system’s motions in a multi-dimensional (mD) reaction-coordinate (RC) space in general; three RCs, denoted as *λ*^(*h*)^(*h* = *α,β,γ*), are introduced for this study. Consequently, “multi-dimensional (mD)” means three-dimensional (3D) for these discussions. The definition of an RC is presented in section 1 of SI. The actual set of the three RCs is given later.

The outline of the method is as follows. The entire conformational space is first divided into many pieces of local spaces as the particular restricted regions defined by the RCs. In every restricted local conformational space, MD simulation is performed independently at an identical constant temperature. This simulation in the local space is called as the Virtual-state coupled MD (VcMD) simulation.^38, 39^ Each local space is overlapped with the neighboring local spaces, and after a particular short period of the MD simulation, the system can transit to the neighboring local spaces depending on the transition probabilities between the overlapped local spaces. The transition probabilities can be set tentatively to appropriate values. Then, after updating the transition probabilities, VcMD simulations are performed in the next short period, and this procedure is continued until resultant probabilities assigned to the local spaces converge to a stationary distribution. Finally, one can calculate the canonical distribution for the entire RC space from the resultant stationary distribution. Because the iterative procedure used in the original mD-VcMD^38, 39^ is time consuming to attain a convergence for a complicated system with many local minima in the free energy landscape, we developed a new procedure named as the GA-guided mD-VcMD, where the Genetic Algorithm is applied to accelerate attaining the stationary distribution even for the complicated system.^40^

The GA-guided mD-VcMD produces a canonical ensemble: a thermally equilibrated conformational ensemble at a simulation temperature. We set the simulation temperature to 300 K for this study and assigned thermodynamic weight *w_i_* to each snapshot, where *i* represents the *i*-th snapshot in the ensemble. Therefore, we can calculate the distribution function of a quantity at equilibrium if the quantity is expressed by the system conformation.

### Systems studied

We examined two systems in this study: pMLF1/14-3-3ε and npMLF1/14-3-3ε systems. The former system consists of 14-3-3ε and pMLF1 (MLF1 phosphorylated at the 34th serine residue: Ace-Arg[31]-Ser[32]-Phe[33]-Sep[34]-Glu[35]-Pro[36]-Phe[37]-Gly[38]-Nme, where Sep represents a phosphorylated serine, and where Ace and Nme are the acetyl and N-methyl groups, respectively); the latter represents 14-3-3ε and npMLF1 (MLF1 non-phosphorylated at the 34th serine residue of Ace-Arg[31]-Ser[32]-Phe[33]-Ser[34]-Glu[35]-Pro[36]-Phe[37]-Gly[38]-Nme). The biomolecules in both systems are surrounded by an explicit solvent (water molecules and ions). Detailed methods used to generate the systems and the system construction are explained in Section 2 of SI.

### Three reaction coordinates

We introduced three RCs to sample various conformations for two systems by GA-guided mD-VcMD. The basic method to define an RC was explained in section 1 of SI. Then, three RCs *λ*^(*α*)^, *λ*^(*β*)^, and *λ*^(*σ*)^ were defined by the inter-mass-center distances between the two atom groups 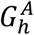 and 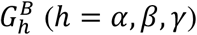, as presented in Figure 1. Details of the atom groups are given in Table S1 of section 4 of SI.

**Figure 1.**
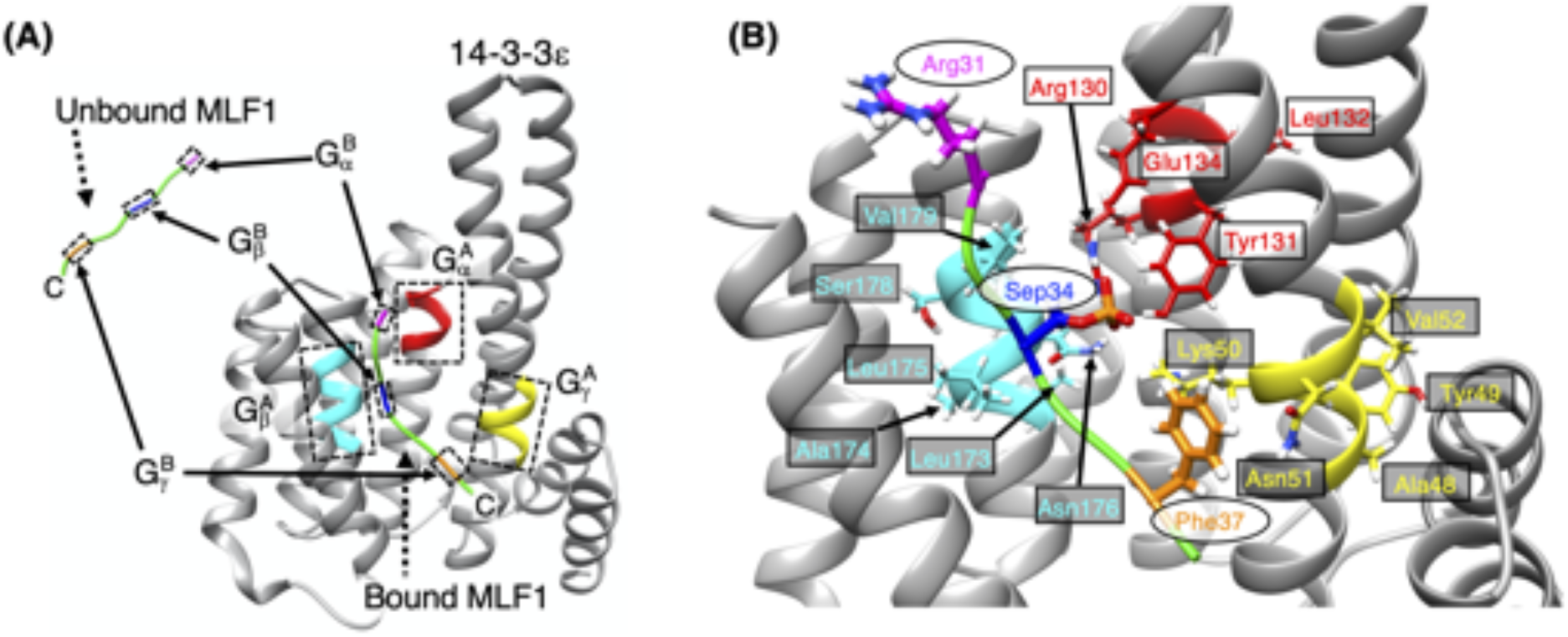
Atom groups to define three RCs *λ*^(*h*)^(*h* = *α,β,γ*) for pMLF1/14-3-3ε. (A) Receptor 14-3-3ε and ligand pMLF1 are shown respectively in gray and green. Bound and unbound forms of pMLF1 are presented. *λ*^(*α*)^ is defined by the mass-center distance between two atom groups 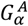 (red fragment) and 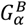 (magenta), *λ*^(*β*)^ by 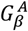 (cyan) and 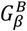 (blue), and *λ*^(*γ*)^ by 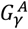 (yellow) and 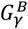 (golden yellow). Each atom group is surrounded by a broken-line rectangle. Section 1 of SI for definition of atom groups. Character “C” is assigned to the C-terminal of pMLF1. (B) Sidechains involved in the atom groups are shown explicitly with the same coloring. Residues in pMLF1 and 14-3-3ε are shown respectively as ovals and rectangles. The npMLF1/14-3-3ε system is not shown. Table S1 of SI presents details of atom groups. Residues Ala 133, Asn 176, and Phe 177 of 14-3-3ε are not shown because they are located at positions behind.

Here, we explain the RCs briefly: *λ*^(*β*)^ represents the distance between Sep 34/Ser 34 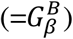 of pMLF1 or npMLF1 and a helical segment 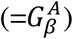 of 14-3-3ε near Sep 34/Ser 34 in the crystal structure. This distance controls the motion of Sep 34/Ser 34 and consequently, the salt-bridge formation between 14-3-3ε and pMLF in the pMLF1/14-3-3ε system, whereas 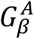 does not involve the counterpart of the salt bridge. For the pMLF1/14-3-3ε system, *λ*^(*β*)^ simply controls the approach of npMLF1 to 14-3-3ε. The RCs *λ*^(*α*)^ and *λ*^(*σ*)^ control the motions of the N-terminal and C-termini of pMLF1 or npMLF1. In general, smaller RCs have closer conformation to the crystal structure. The RCs for the crystal structure are *λ*^(*α*)^ = 15.6 Å, *λ*^(*β*)^ = 7.9 Å, and *λ*^(*σ*)^ = 8.7 Å.

### Initial conformations of simulation

The initial conformation of the GA-guided VcMD simulation is not the crystal complex structure but one for which pMLF1 (or npMLF1) is distant from the position in the native complex structure. The conformation of pMLF1 (or npMLF1) is randomized. We generated those conformations according to the method explained in section 3 of SI. Figure S5 presents additional details. We generated 256 initial conformations because we performed 256 runs of GA-guided VcMD as described below.

### Simulations

Although the current system is a single heterodimer (Figure S4C), 14-3-3ε maintained its tertiary structure in preliminary NVT simulations (data not shown). Therefore, no structural restraint was introduced to maintain the 14-3-3ε structure during the GA-guided mD-VcMD simulations. To raise the sampling efficiency, we performed 256 runs in parallel^41,42^ for both system starting from randomized conformations generated in the section above and in section 3 of SI.

The GA-guided mD-VcMD simulation was performed using a computer program, omegagene/myPresto,^43^ with the following condition: The SHAKE algorithm^44^ is used to fix the covalent-bond lengths related to hydrogen atoms, the Berendsen thermostat to control temperature,^45^ the zero-dipole summation method^46–48^ for long-range electrostatic computations, time-step of 2 fs (Δ*t* = 2 fs), and simulation temperature of 300 K. An ensemble from the Berendsen thermostat can approximate a canonical distribution for a system of many atoms, whereas it generates a non-physical distribution for a small system.^49^ To compute the original potential energy of the system, the Amber hybrid force fields^50^ (mixture parameter *w* = 0.75) were used for pMLF1, npMLF1, and 14-3-3ε, the TIP3P model for water molecule,^51^ and the Joung–Cheatham model for chloride and sodium ions.^52^

## RESULTS

### Conformational distribution in 3D-RC space

We show in this section that the two systems provide greatly different conformational distributions from one another. The most stable state for pMLF1/14-3-3ε involved crystal structure-like conformations. That for npMLF1/14-3-3ε consists of conformations far from the crystal structure.

We performed GA-guided mD-VcMD simulations and obtained an equilibrated conformational ensemble at 300 K for each system. The simulation condition is shown in section 4 of SI. We calculated a 3D density function, denoted as *ρ*_3*D–RC*_(*λ*) = *ρ*_3*D–RC*_(*λ*^(*α*)^,*λ*^(*β*)^, *λ*^(*γ*)^, in the 3D-RC space from the obtained ensemble: The probability of detecting the system in a small RC volume *dλ* = *dλ*^(*α*)^*dλ*^(*β*)^*dλ*^(*γ*)^ is given as *ρ*_3*D–RC*_(*λ*)*dλ*. Figure 2 shows the resultant density functions for pMLF1/14-3-3ε (A) and npMLF1/14-3-3ε (B). The highest density region (i.e., the lowest free-energy region or the most thermodynamically stable region) differed between the two systems. In pMLF1/14-3-3ε, as shown later, the most stable region involved complex structures similar to the crystal structure. In npMLF1/14-3-3ε, the most stable region consists of conformations where npMLF1 and 14-3-3ε were unbound or were slightly contacting. Later, we demonstrate that this difference in the most stable region derives from the presence or absence of salt-bridges between MLF1 and 14-3-3ε. Accuracy and convergence of distribution *ρ*_3*D–RC*_(*λ*) for both systems are discussed in Figure S6 of SI.

**Figure 2.**
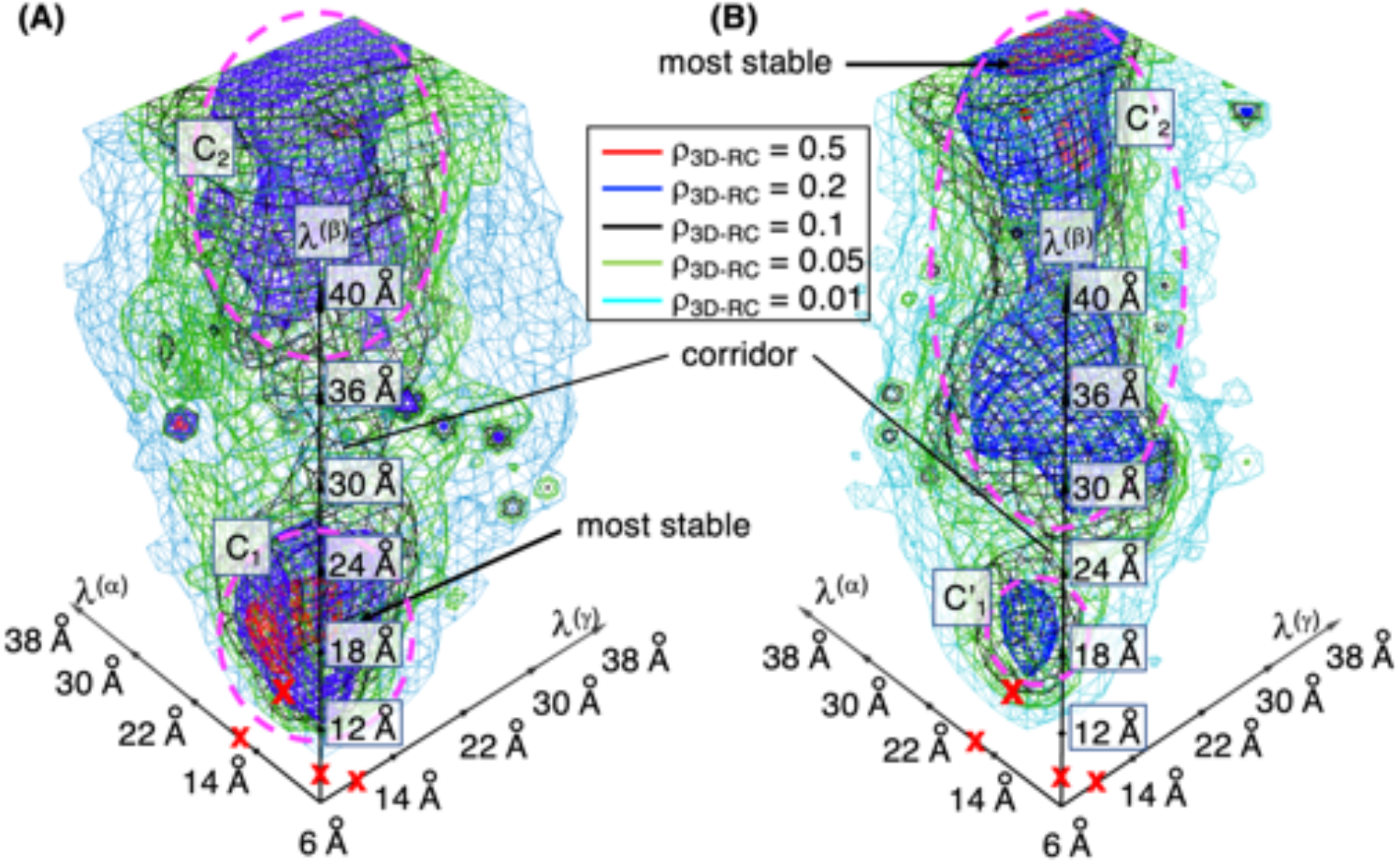
Distribution *ρ*_3*D–RC*_(*λ*^(*α*)^,*λ*^(*β*)^,*λ*^(*λ*)^) of (A) pMLF1/14-3-3ε and (b) npMLF1/14-3-3ε systems in 3D-RC space, which is calculated from equilibrated conformational ensemble at 300 K for each system. Contour levels of the distribution are shown in inset. The distribution is normalized as ∫ *ρ*_3*D*–*RC*_(*λ*^(*α*)^, *λ*^(*β*)^, *λ*^(*γ*)^ *dλ*^(*α*)^ *dλ*^(*β*)^ *dλ*^(*γ*)^ = 1. Clusters are shown by broken-line circles with labels *C*_1_, *C*_2_, 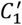 and 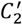. Corridors connecting *C*_1_ and *C*_2_ as well as 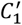 and 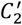 are indicated by “corridor”. Positions of the most stable (highest density) regions are indicated by “most stable”. The RCs for the crystal structure are indicated by red-colored crosses on their axes: *λ*^(*α*)^ = 15.6 Å, *λ*^(*β*)^ = 7.9 Å, and *λ*^(*γ*)^ = 8.7 Å. Red-colored cross in the distribution indicates the crystal-structure position.

Whereas the two systems provided different landscapes, two clusters existed in common. We designated them as *C*_1_ and *C*_2_ for pMLF1/14-3-3ε and 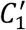 and 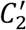 for npMLF1/14-3-3ε as in Figure 2. Clusters *C*_1_ and *C*_2_ correspond roughly to 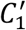 and 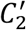, respectively. In either system, the two clusters are connected by a corridor. Cluster *C*_1_ is more stable than *C*_2_, whereas 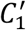 is considerably less stable
 than 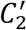. Cluster 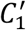 exists marginally as a cluster.

Here, to assess the conformational difference of pMLF1 (or npMLF1) between a snapshot and the crystal structure, we defined a measure *RMSD_lig_*: the mainchain heavy-atomic root mean square deviation for residues from Phe 33 to Phe 37 in pMLF1 (or npMLF1) after superposition of the 14-3-3ε protein part between the snapshot and the crystal structure. Therefore, *RMSD_lig_* reflects not only the internal motions of pMLF1 (or npMLF1) but also its translational and rotational motions relative to 14-3-3ε. As described in the *Introduction*, the contribution of aminoacids to the complex stability was measured for Phe 33, Glu 35, and Phe33. The most important residue for the complex formation is Sep 34.^7^ Consequently, these amino-acids exist in residues 33–37. Residues Phe 33 and Phe 37 are located at the edges for the residue region.

Figures 3A and 3B present conformations of the pMLF1/14-3-3ε system taken from cluster *C*_1_. Panel A displays the pMLF1 conformation with the smallest *RMSD_lig_* (0.69 Å). The sidechain positions of Sep 34 and Phe 37 coincide well with those in the crystal structure shown in yellow. Although Phe 33 is deviated from the crystal-structure position, this residue interacts with Leu 230 via a hydrophobic contact, as discussed later. Panel B is a conformation picked randomly from cluster *C*_1_ (*RMSD_lig_* = 4.0 Å). Whereas Phe 33 and Phe 37 were deviated from the crystal-structure positions, the position of Sep 34 agreed well with the crystal-structure position. We present three conformations picked randomly from cluster *C*_2_ in Figure S7A of SI, where pMLF1 was far from the crystal-structure position, and contacting to 14-3-3ε (encounter complexes) or isolated in solution.

**Figure 3.**
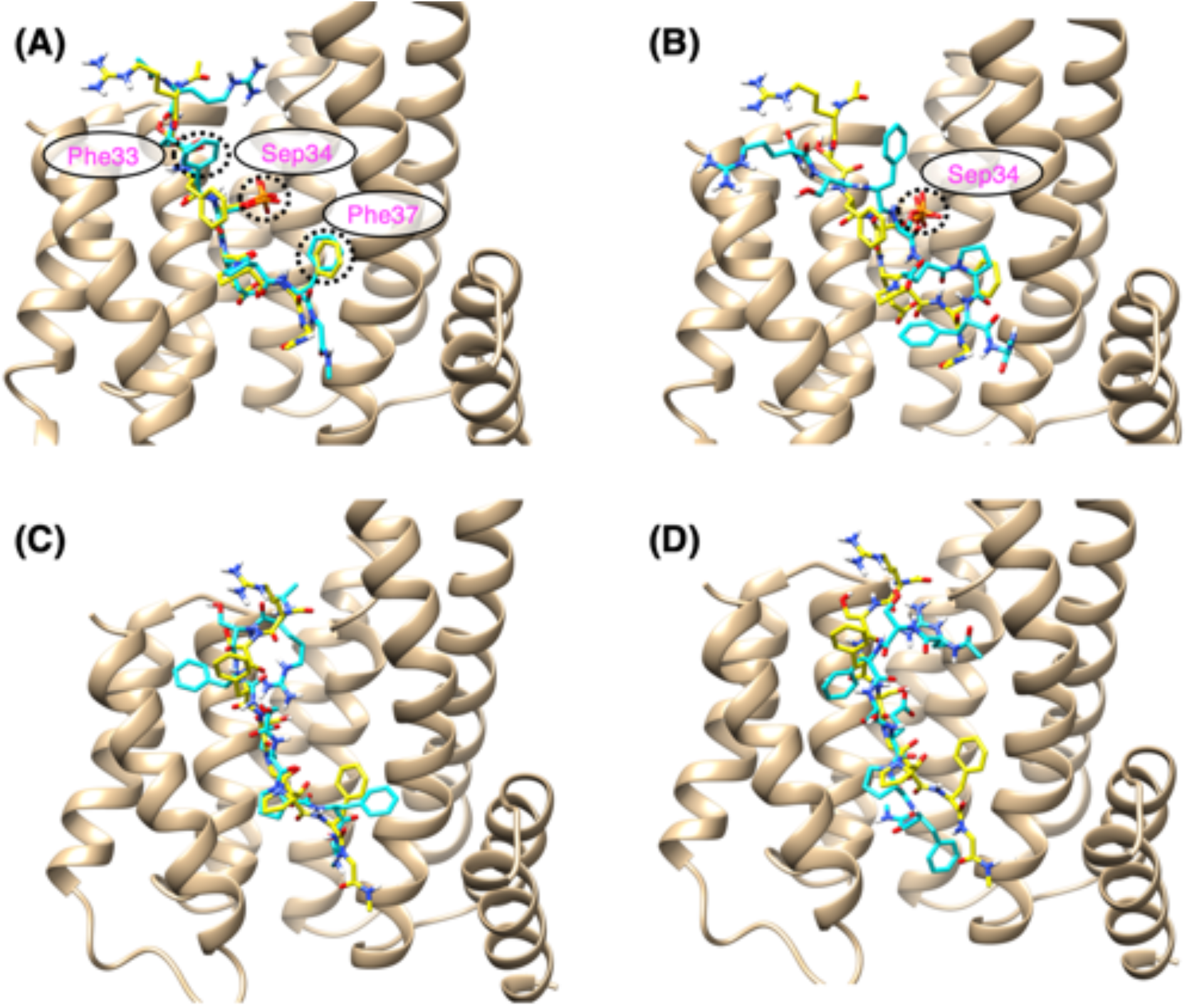
Panels (A) and (B) display conformations from the pMLF1/14-3-3ε system; (C) and (D) show those from the npMLF1/14-3-3ε system. Yellow-colored MLF1 conformations in all panels are those from the crystal structure. Residues denoted by ovals belong to pMLF1. The pMLF1 in (A) is the smallest *RMSD_lig_* conformation from cluster *C*_1_. The pMLF1 in (B) is a randomly chosen conformation from cluster *C*_1_. Residues Phe 33, Sep 34, and Phe 37 in (A) and Sep 34 in (B) are indicated by broken-line circles. Panel (C) is the smallest *RMSD_lig_* conformation from cluster 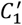. Panel (D) is one randomly chosen conformation from cluster 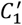.

Figures 3C and 3D display conformations of the npMLF1/14-3-3ε system from 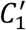. Panel C is the smallest *RMSD_lig_* conformation in cluster 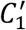 (*RMSD_lig_* = 1.60 Å). Panel D presents a conformation chosen randomly from cluster 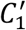 (*RMSD_lig_* = 3.3 Å). Three conformations selected randomly from cluster 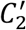 are shown in Figure S7B of SI.

### Distribution of MLF1 around 14-3-3ε at 300 K

The earlier section showed that a single-residue difference (Sep/Ser) of MLF1 greatly varied the conformational distribution. Consequently, phosphorylation of MLF1 crucially governs the complex stability. This section presents further analysis of the difference of the complex stability between the two systems by calculating one-dimensional radial distribution functions.

As described in the *Introduction,* Molzan *et al.* reported that Phe 33 contributed greatly to complex-structure stabilization, in contrast to Glu 35 and Phe 37, which did not contribute substantially to the stability.^7^ Therefore, we specifically examined the following four inter-residue distances: The first distance *r*^*Phe*33^ is the minimum distance from the ring center of Phe 33 of pMLF1 (or npMLF1) to the sidechain heavy atoms of the hydrophobic interaction partners, Leu 223 and Leu 230, of 14-3-3ε (Figure S2). The second distance *r*^*Sep*34^ (or *r*^*Ser*34^) is the minimum distance from the Og atom of Sep 34 of pMLF1 (or Ser 34 of npMLF1) to the Cζ atoms of Arg 57 and Arg 130 of 14-3-3ε. The positions of these three residues are presented in Figure S8 of SI. We did not select the phosphate atom for Sep 34 for the definition of *r*^*Sep*34^ because Ser 34 does not involve a phosphate atom. To compare behaviors of Sep 34 and Ser 34, the same atom should be selected. The third distance *r*^*Glu*35^ the minimum distance from the Cγ atoms of Glu 35 of pMLF1 (or npMLF1) to the N*ζ* atoms of the salt-bridge partner, Lys 50 and Lys 123, of 14-3-3ε. The fourth distance *r*^*Phe*37^ the minimum distance from the ring center of Phe 37 of pMLF1 (or npMLF1) to the heavy atoms in the stem of Lys 50 of 14-3-3ε. A lysine sidechain has a long hydrophobic stem.

We calculated the radial distribution function, *p*(*r*^*μ*^) with respect to the four distances defined above (*μ* = *Phe*33, *Sep*34/*Ser* 34, *Glu*35, or *Phe*37). Details for the calculation of *p*(*r*^*μ*^) are presented in section 5 of SI. Figure 4 depicts the calculated radial distribution functions. A remarkable difference between the two systems is related to Sep/Ser 34: the sharp peak of *p*(*r*^*Sep*34^) is portrayed in Figure 4A, although *p*(*r*^*Ser*34^) shows broad peaks (Figure 4B). This difference is attributable to the presence or absence of the salt bridges associated with Sep 34 and Ser 34, as discussed later. According to the decrease of the peak height from *p*(*r*^*Sep*34^) to *p*(*r*^*Ser*34^), the peaks of the other radial distribution functions decreased. This result derives naturally from Figure 3B: Cluster 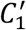 is less stable than 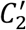 in the npMLF1/14-3-3ε system.

**Figure 4.**
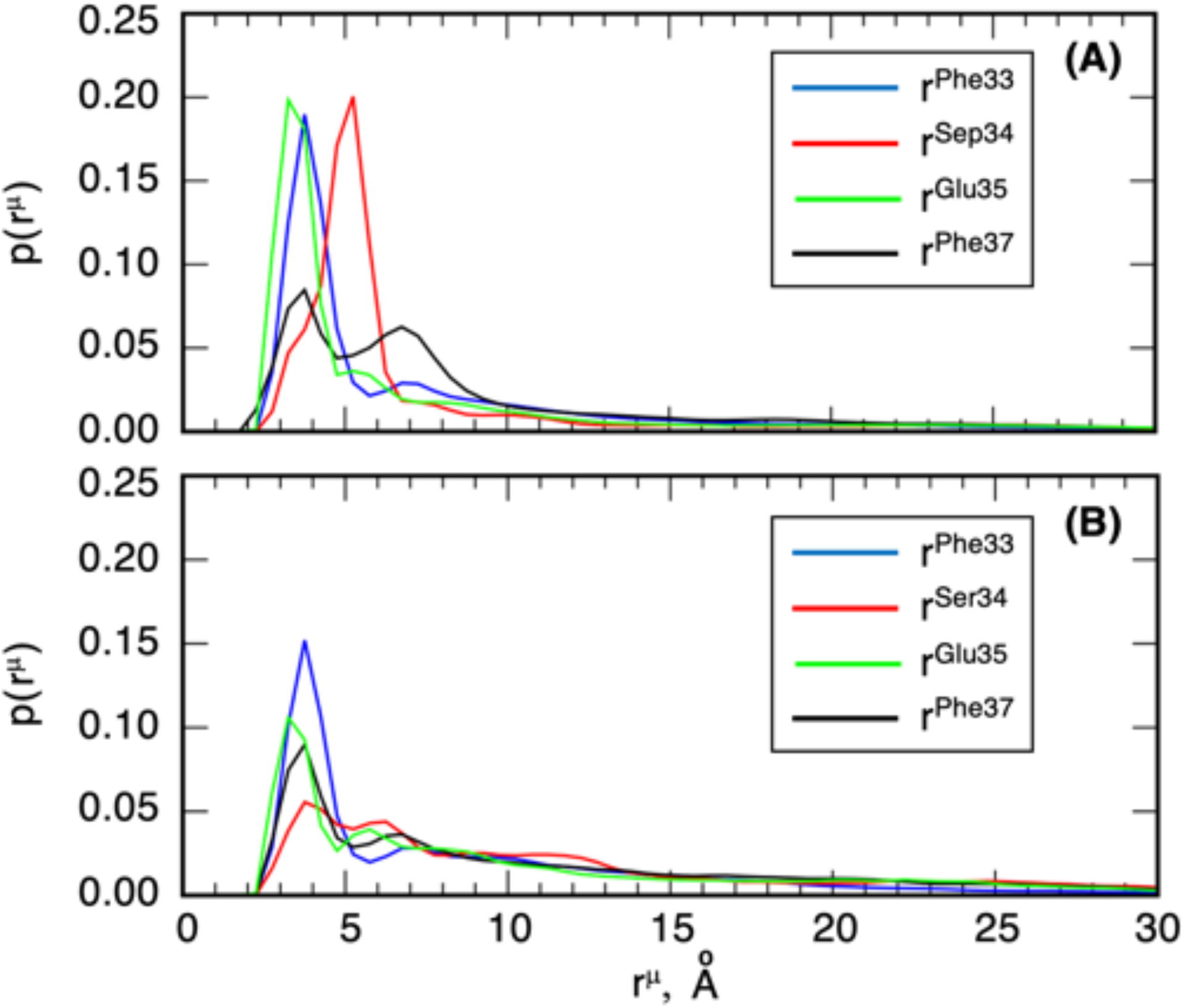
Radial distribution function *p*(*r*^*μ*^) for four distances: *μ* = Phe33, Sep/Ser34, Glu35, and Phe37. See section 5 of SI for definition of *p*(*r*^*μ*^). Panels (A) and (B) respectively show pMLF1/14-3-3ε and npMLF1/14-3-3ε.

Furthermore, in the pMLF1/14-3-3ε system, it is noteworthy that *p*(*r*^*Phe*33^) has a sharp peak, whereas the distribution of *p*(*r*^*Phe*37^) is broad (Figure 4A). This result supports the experimental data showing that Phe 33 of pMLF1 contributes more to complex stability than Phe 37 of pMLF1 does.^7^ Later, this point is discussed further.

The situation of Glu 35 of pMLF1 is problematic: *p*(*r*^*Glu*35^) has a sharp peak at *r*^*Glu*35^ ≈ 3 Å in the pMLF1/14-3-3ε system (Figure 4A). Consequently, Glu 35 seems to form a salt bridge to Lys 50 or Lys 123 of 14-3-3ε in the simulation. This result might indicate that Glu 35 contributes to the complex stability. However, as described in the *Introduction,* the mutation E35A of pMLF1 did not change the complex stability. Later, we analyze fluctuations of Glu 35 around 14-3-3ε to explain this discrepancy rationally.

One might note that most plots of Figure 4 had multiple peaks (or shoulders). The origin of this multiple-peak feature is the existence of multiple conformational clusters of MLF1 on the 14-3-3ε surface, as shown later by 3D spatial distributions.

### 3D spatial distribution of MLF1 around 14-3-3ε at 300 K

We further investigate the 3D-spatial distribution (density map), 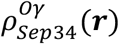 and 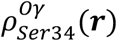, of the Sep 34 and Ser 34 positions around 14-3-3ε to view the shape of the free-energy landscape for the positions of Sep 34 and Ser 34. The method used to compute 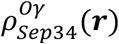 and 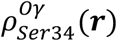 is given in section 6 of SI. In fact, the distribution can be converted to the potential mean force (PMF) simply as 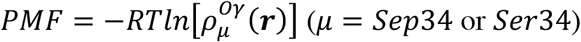.

Figure 5A shows density maps of 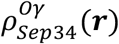, which is the spatial density of the O*γ* atom of Sep 34 of pMLF1. Apparently, in the pMLF1/14-3-3ε system, the high-density spot was found around the O*γ*-atomic position of Sep 34 in the crystal structure (red contours in Figure 5A). This figure also shows that Sep 34 is distributed over almost the entire surface of 14-3-3ε at a lower density level (dark green contours). However, at a middle density level (magenta-colored contours), the distribution was inhomogeneous with four large clusters *D*_1_, …, *D*_4_. Cluster *D*_1_ involved the high-density spot described above. The other clusters did not involve such a high-density region. Consequently, *D*_1_ was the most stable of the four clusters thermodynamically. Clusters *D*_1_ and *D_2_* were connected smoothly without density discontinuity: In other words, *D*_1_ and *D_2_* might be a single cluster. In contrast, density discontinuity (free-energy barriers) was found among clusters *D*_1_, *D*_3_, and *D*_4_.

**Figure 5.**
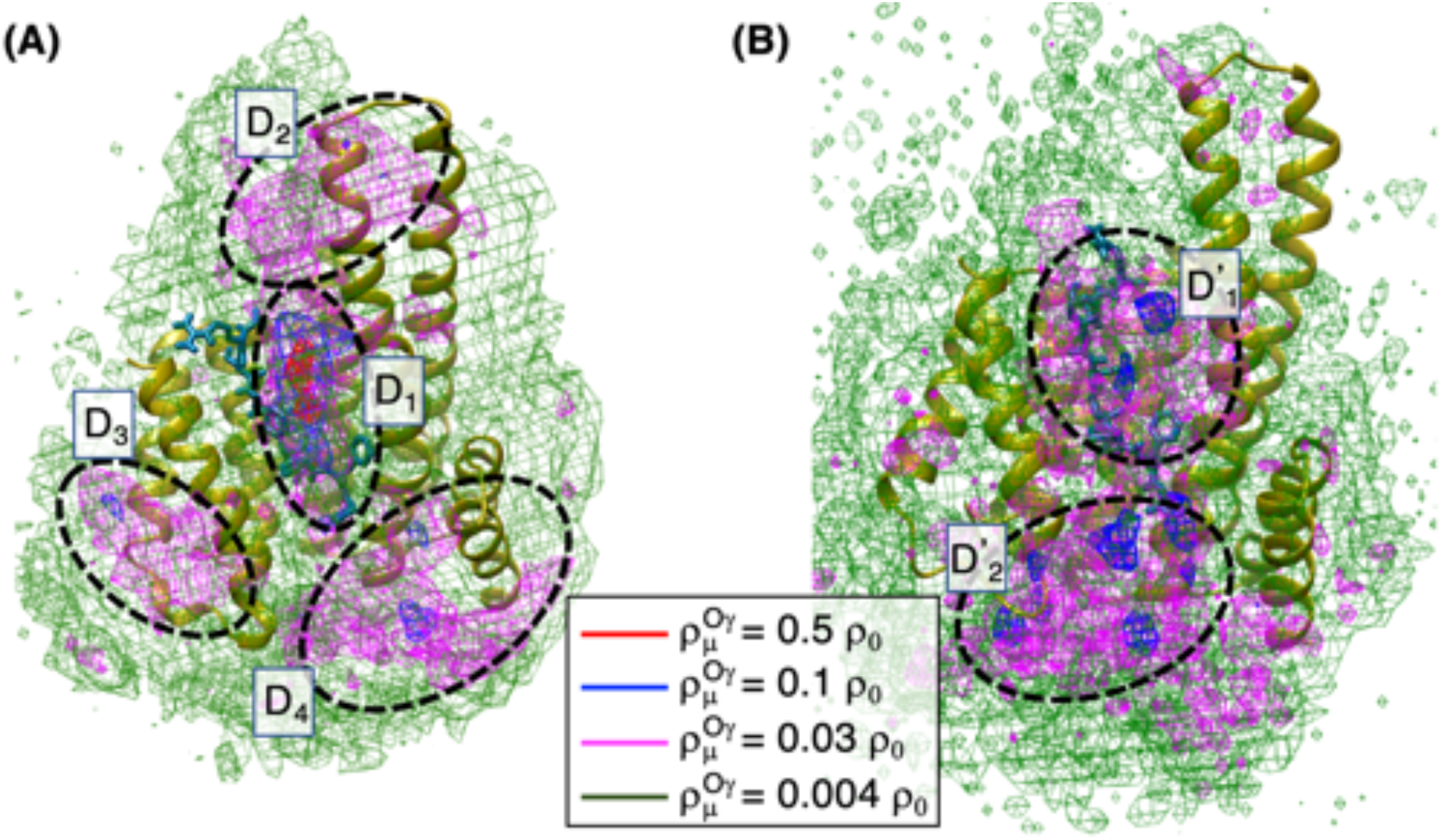
Density contour map 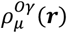 of the O*γ* atomic position of residue 34 of MLF1: (A) 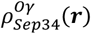 and (b) 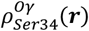. Contour levels are shown in the inset, where *ρ*_0_ = 0.01. The molecular structures shown are the crystal complex structure. Clusters surrounded by broken-line circles, designated as *D_i_* (*i* = 1, …,4) and 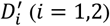, are described in text.

Figure 5 shows that GA-guided mD-VcMD can sample not only high-density regions but also low-density ones. Therefore, we presume that oversight of low-probability conformations did not occur. The minor-basin sampling might be fundamentally important to investigate connections among major basins, as described in the *Introduction*.

To analyze cluster *C*_1_ (Figure 2A), we calculated the density map of 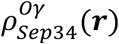 using snapshots only from cluster *C*_1_. Figure S9 of SI shows that conformations in *C*_1_ involved cluster *D*_1_ and a part of *D*_2_ of Figure 5A. It is particularly interesting that clusters *D*_3_ and *D*_4_ are not found in Figure S9. Remember that clusters *D*_3_ and *D*_4_ are not interesting for discussion of binding pathways because free-energy barriers exist between *D*_3_ and *D*_1_ and between *D*_4_ and *D*_1_. However, the existence of *D*_3_ and *D*_4_ is important to characterize the free-energy landscape of this system: This landscape is not funnel-like but rather multi-modal, as found computationally before.^37,53^

Figure 5B is a contour map of 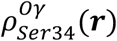 from the npMLF1/14-3-3ε system. No red contours were found in this density map, which results from absence of a salt bridge at the 34th residue. We found two clusters, denoted as 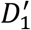 and 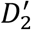, in Figure 5B at the level of 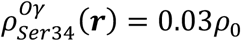 (magenta-colored contours). The position of 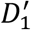 overlaps that of 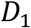 in Figure 5A, revealing that npMLF1 has a weak tendency of staying around the pMLF1 position in the crystal structure.

### Salt-bridge relay between pMLF1 and 14-3-3ε

As presented in Figure 5A, cluster *D*_1_ was the most stable of the four clusters. Here, we investigate the internal structure of cluster *D*_1_. Figure 6A shows that the contours of 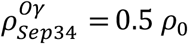 are split into two spots, named *E*_1_ and *E*_2_, and that *E*_1_ involves the crystal-structure position of the O*γ* atom of Sep 34 of pMLF1.

**Figure 6.**
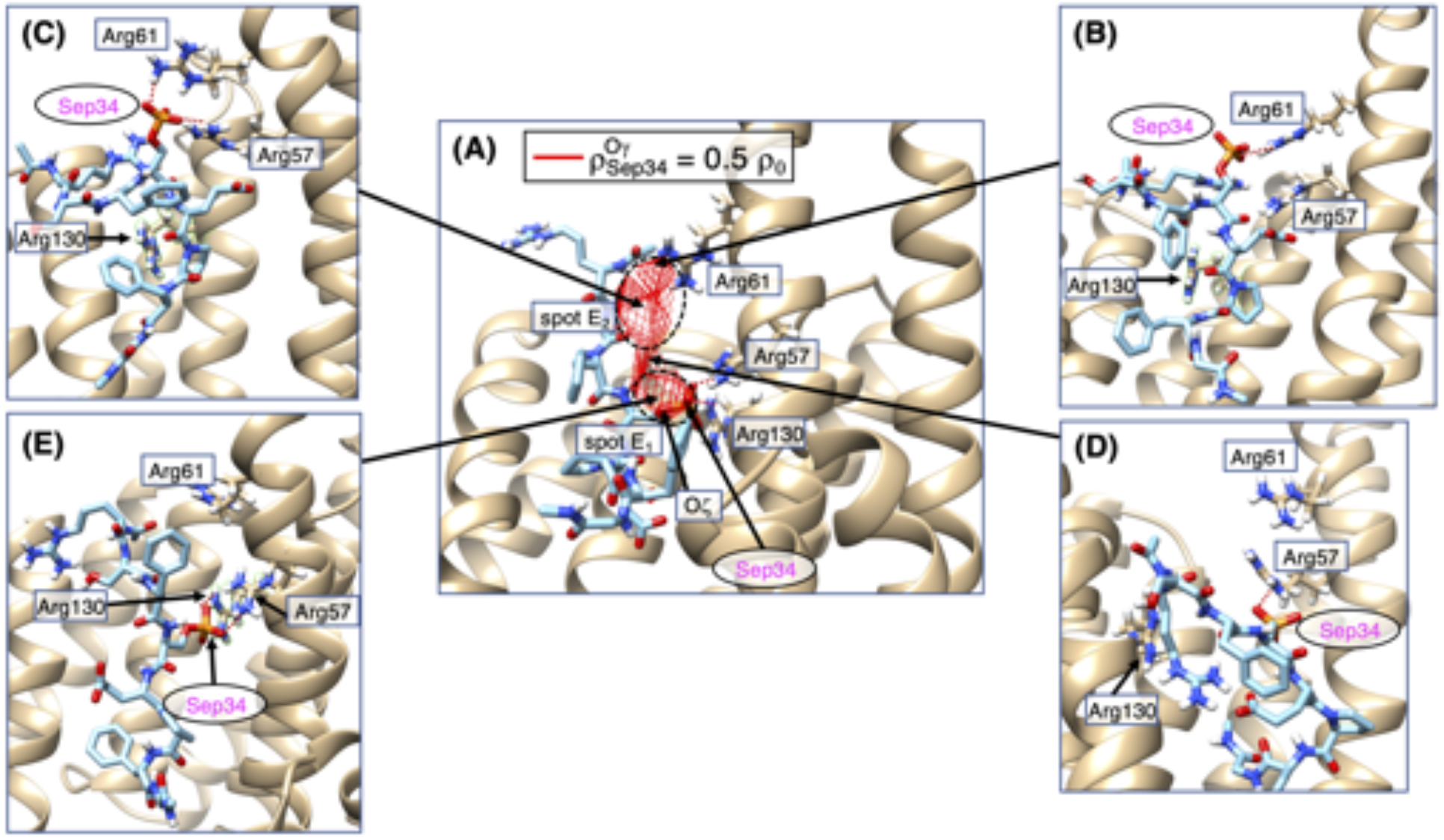
(A) Close-up of cluster *D*_1_. Red-colored contours are those with 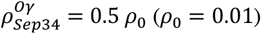. Two high-density spots *E*_1_ and *E*_2_ are shown. (B) Snapshot taken from the fringe of *E*_2_. (C) Snapshot taken from the middle of *E*_2_. (D) Snapshot taken from the corridor connecting *E*_1_ and *E*_2_. (E) Snapshot taken from *E*_1_. The salt bridge is shown as a brown broken line. Residues labeled in ovals belong to pMLF1, and those in rectangles do to 14-3-3ε.

As described above, a probable binding pathway is that from cluster *D*_2_ to *D*_1_ because no free-energy barrier is apparent between *D*_1_ and *D*_2_. Therefore, the conformational change of pMLF1 from *D_2_* to *D*_1_ is noteworthy. Figure 6B is a snapshot taken from a fringe of *E*_2_, where Sep 34 formed a salt bridge to Arg 61 of 14-3-3ε, which is not observed in the crystal structure. Figure 6C displays a snapshot taken from the middle of *E*_2_, where Sep 34 formed salt bridges to Arg 61 and Arg 57 of 14-3-3ε. The salt bridge to Arg 57 is one observed in the crystal structure. Additionally, it is noteworthy that Arg 130 of 14-3-3ε did not participate in the salt bridge to Sep 34. Figure 6D portrays a snapshot selected from the corridor between *E*_1_ and *E*_2_, which shows that the salt bridge to Arg 61 was broken and that the salt bridge to Arg 130 was not formed. Consequently, the conformation at the corridor had only one salt bridge, which suggests that the conformation at the corridor is energetically less stable than that in *E*_2_. Figure 6E presents a snapshot taken from *E*_1_, in which the two salt bridges were formed as in the crystal structure. We presume that the corridor between *E*_1_ and *E*_2_ originated from decrease of a salt bridge. Consequently, Figures 6B–6E suggest that the salt-bridge relay takes place when the conformation moves from *E*_2_ to *E*_1_.

### Spatial distribution of Phe 33 and Phe 37 of MKF1 around 14-3-3ε at 300 K

As described in the *Introduction*,^7^ Phe 33 of pMLF1 contributes greatly to the complex stability, whereas Phe 37 does not. Next, we investigate the spatial distributions of Phe 33 and Phe 37 of pMLF1 around 14-3-3ε to elucidate the contributions of these residues to the complex stability.

Figure S2 displays the pMLF1/14-3-3ε complex structure (crystal structure), where Phe 33 of pMLF1 is sandwiched between Leu 223 and Leu 230 of 14-3-3ε, and Phe 37 of pMLF1 contacts to the hydrophobic sidechain stem of Lys 50 of 14-3-3ε. We calculated the density of the sidechain ring centers of Phe 33 and Phe 37, which are expressed respectively by 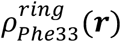 and 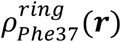 of pMLF1 (Figure 7A) or npMLF1 (Figure 7B). Section 6 of SI presents the procedure used to calculate the densities. For the pMLF1/14-3-3ε system, Phe 33 distributed in a region named spot *H* in Figure 7A, which is capable of contacting Leu 223 and/or Leu 230. Spot *H* is the largest spot for Phe 33. It is the origin of the sharp peak of *r*^*Phe*33^ in Figure 4A.

**Figure 7.**
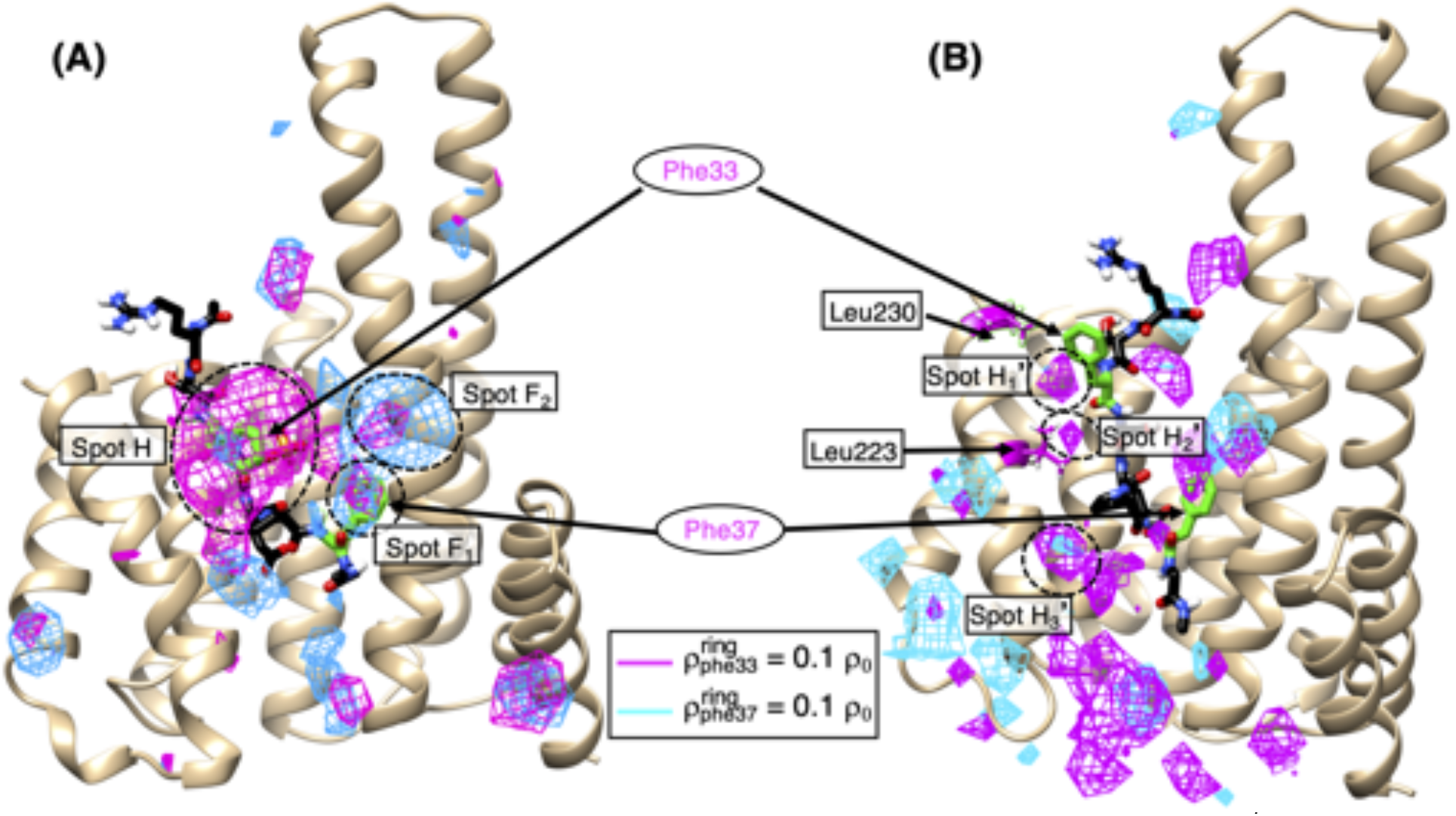
Spatial distributions of the sidechain ring-center positions of Phe 33 and Phe 37, denoted respectively as 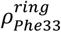 (magentacolored contours) and 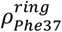 (cyan-colored contours). The contour level is 0.1 *ρ*_0_ (*ρ*_0_ = 0.01) for both residues. Panels (A) and (B) respectively show pMLF1/14-3-3ε and npMLF1/14-3-3ε. The displayed molecular structures are the crystal structure, in which green residues are Phe 33 and Phe 37. Residues labeled in ovals and rectangles respectively belong to MLF1 and 14-3-3ε. In panel (A), broken-line circles named spots *H*, *F*_1_, and *F*_2_ are described in the text. In panel (B), Leu 223 and Leu 230 are shown in magenta color. Three broken-line circles, named spots 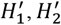 and 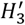, are described in the text.

Two spots *F*_1_ and *F*_2_ for 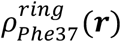 in Figure 7A are related to the two peaks of 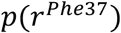 in Figure 4A. That is, *F*_1_ and *F*_2_ respectively represent the origins of the peaks at *r*^*Phe*37^ ≈ 4 Å and *r*^*Phe*37^ ≈ 7 Å. In contrast to the large spot *H* for Phe 33, both spots *F*_1_ and *F*_2_ for Phe 37 were small, which suggests that Phe 37 contributes less to stabilization of the complex structure.

In Figure 7B, the red and blue contours distributed widely and sparsely around 14-3-3ε. It is noteworthy that Ser 34 in npMLF1/14-3-3ε distributed in clusters 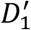 and 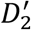 in Figure 5B. Figures 5B and 7B show that npMLF1 has no strong preferential spot. Although the three broken-line circles, named as spots 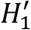, 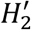, and 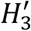 in Figure 7B, were found to be close to Leu 223 and Leu 230, these spots were small. They caused the peak of *r*^Phe*33*^ in Figure 4B.

We calculated the 3D distributions 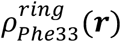 and 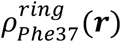 for the pMLF1/14-3-3ε system using snapshots only from cluster *C*_1_ in Figure 2A. They are portrayed in Figure S10 of SI. Figure is similar to Figure 7A: 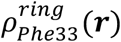 is localized around the position of Phe 33 in the crystal structure; also, 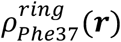 distributed broadly on the surface of 14-3-3ε. However, the density level of Figure S10 was higher than that of Figure 7A for both Phe 33 and Phe 37 because the distribution was normalized only using snapshots in *C*_1_ in Figure S10.

### Spatial distribution of Glu 35

In the crystal structure, the sidechain of Glu 35 of pMLF1 forms salt-bridges to Lys 50 and Lys 123 of 14-3-3ε. However, as described in the *Introduction,* Glu 35 of pMLF1 does not contribute to the complex stability.^7^ To explain this discrepancy, we calculated the density maps 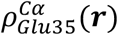 and 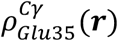, respectively, for the Cα and Cγ atomic positions of Glu 35 of pMLF1 for the pMLF1/14-3-3ε system. Section 6 of SI explains the procedure used to calculate the densities.

Figure 8A shows high-density regions drawn around the crystal structure, in which Glu 35 forms salt bridges to Lys 50 and Lys 123 of 14-3-3ε. The contours show that both the Cα and Cγ atoms shifted from the crystal-structure positions toward the solvent, and that the mainchain of Glu 35 fluctuated to a considerable degree, as represented by the broken-line arrow along the magenta-colored contours in Figure 8A. We consider that this positional shift hinders Glu 35 of pMLF1 to form salt bridges to Lys 50 and Lys 123 of 14-3-3ε. The black contours in Figure S11A demonstrate that Glu 35 can take a position (broken-line circle of Figure 11A), from which Glu 35 was able to form the salt bridges to both Lys 50 and Lys 123. However, the density assigned to this position was small 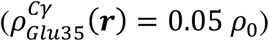. Consequently, it seems that Figure 8A and Figure 4A are mutually contradictory.

**Figure 8.**
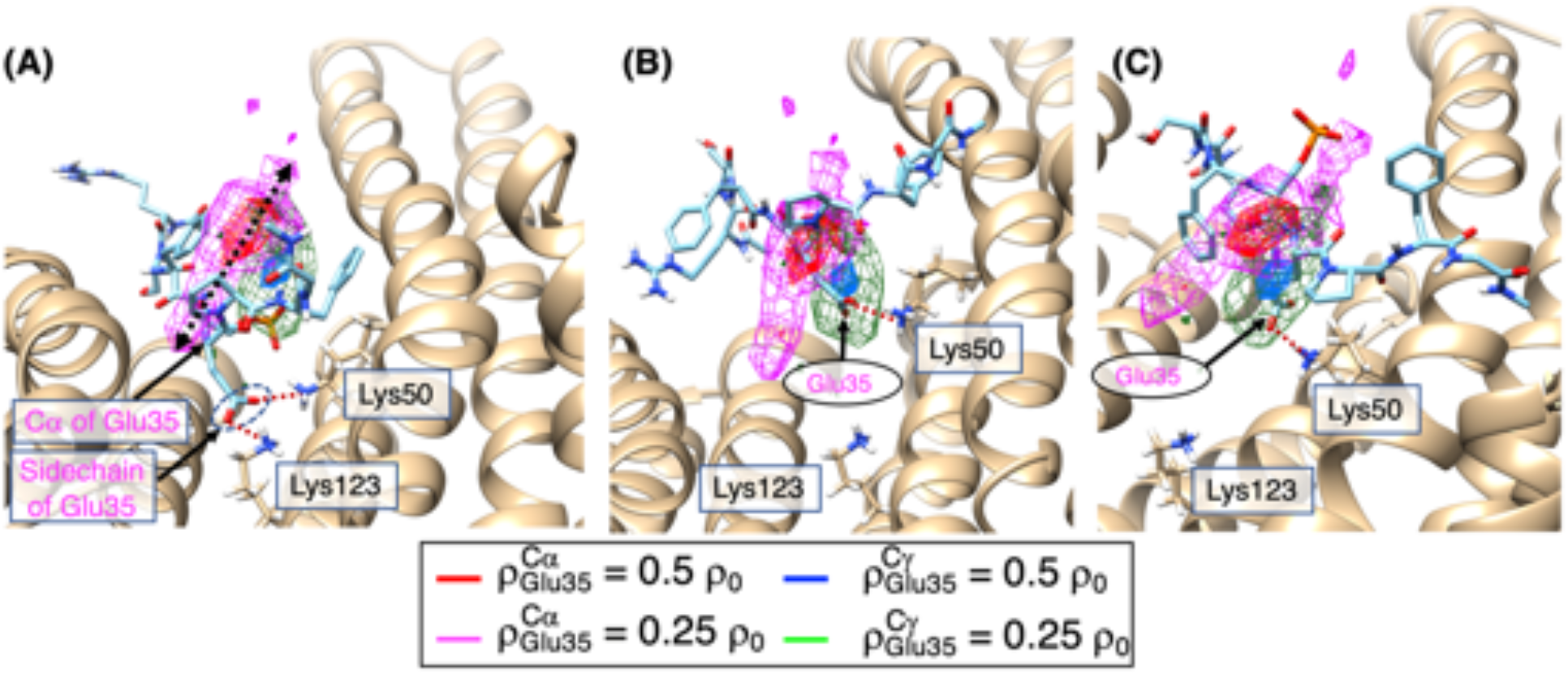
(A) Density contour maps of 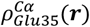 and 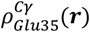 for the pMLF1/14-3-3ε system. The presented molecular structure is the crystal one, where the Cα atom and sidechain tip of Glu 35 of pMLF1 are denoted respectively as “Cα of Glu35” and “Sidechain of Glu35”. Contour levels are shown in the inset (*ρ*_0_ = 0.01). Panels (B) and (C) display snapshots chosen randomly from conformations that are involved in the contour regions of panel (A). Panels (B) and (C) are drawn with different orientations from panel (A) to show Glu 35 clearly. Salt bridge is shown by the brown broken line. Residues labeled in ovals belong to pMLF1. Those in rectangles belong to 14-3-3ε.

To elucidate this discrepancy, we checked snapshots for which Cα and Cγ atoms were involved in the contours of 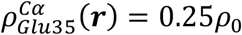 (magenta contours in Figure 8A) and 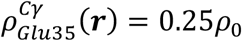 (green contours of Figure 8A), and found that most of those snapshots in the contour regions formed a salt bridge only to Lys 50 of 14-3-3ε. Figures 8B and 8C present two snapshots selected randomly from the contour regions. To clarify this observation, we calculated *p*(*r*^*Glu*35–*Lys*50^) and *p*(*r*^*Glu*35–*Lys*123^): The former is a radial distribution function for the distance *r*^*Glu*35–*Lys*123^ from the Cγ atom of Glu 35 of pMLF1 to the Nζ atom of Lys 50 of 14-3-3ε. The latter is a radial distribution function for the distance *r*^*Glu*35–*Lys*123^ from the Cγ atom of Glu 35 to the N*ζ* atom of Lys 123 of 14-3-3ε. These distances were used to define *r*^*Glu*35^ in Figure 4. Figure S11B clarifies that the salt bridge between Glu 35 and Lys 123 was rarely formed. Furthermore, we calculated salt-bridge-formation probabilities *Q*_*Glu*35–*Lys*50_ and *Q*_*Glu*35–*Lys*123_ according to section 7 of SI. The resultant fraction *Q*_*Glu*35–*Lys*123_ /*Q*_*Glu*35–*Lys*123_ was 0.18 × 10^−5^, which again indicated that the salt bridge between Glu 35 and Lys 50 was dominant.

In fact, Lys 50 of 14-3-3ε is located on the surface of 14-3-3ε. Consequently, the sidechain stem of Lys 50 can modulate its sidechain-tip position to maintain the salt bridge to Glu 35 of pMLF1. In contrast, Lys 123 is somewhat buried in 14-3-3ε. Therefore, the sidechain tip of Lys 123 cannot move together with the positional shift of Glu 35. Consequently, the salt bridge between Glu 35 of pMLF1 and Lys 123 of 14-3-3ε is not formed.

We presume that the positional shift and large fluctuations of Glu 35 of pMLF1 weaken the contribution of Glu 35 to the complex-structure stability. It is noteworthy that the sidechain stem of Lys 50 of 14-3-3ε is the hydrophobic interaction partner of Phe 37 in the crystal structure. Therefore, the positional fluctuations of Glu 35 of pMLF1 affect the sidechain motions of Lys 50 of 14-3-3ε. They therefore hinder the hydrophobic interaction between Phe 37 of pMLF1 and Lys 50 of 14-3-3ε: the small contributions of Glu 35 and Phe 37 to the complex structure stability might be mutually correlated.

### Orientation and ligand conformational variety of pMLF1 around 14-3-3ε

This section presents specific examination of the molecular orientation and the conformational variety of pMLF1 around 14-3-3ε to elucidate the conformational convergence of pMLF1 with approach of the binding site of 14-3-3ε. First, we defined the orientation of pMLF1 by a unit vector ***e**_i_* (|***e***_*i*_| = 1) for snapshot *i,* which is pointing from the Cα atom of Phe 33 to the Cα atom of Phe 37 of the snapshot. The orientation vector ***e**_ref_* (|***e**_ref_*| = 1) for the crystal structure was also calculated similarly.

The ligand pMLF1, for which O*γ* atom of Sep 34 was visiting a cube ***r***, has an orientational variety. We calculated 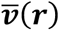, the average of the orientation vectors ***e**_i_* in a cube ***r***, using a method explained in section 8 of SI (Eq. S1). Note that the vector 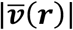 shows a degree of orientational ordering of pMLF1 detected in cube ***r***; also, 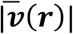 takes the maximum of 1 when all ***e**_i_* have exactly the same orientations in the cube. If snapshots have uncorrelated orientations, then |v(r)| becomes small.

Next, to elucidate the tendency of 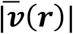 to align to ***e**_ref_* for the crystal structure or not, we calculated a scalar product of 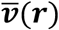 and ***e**_ref_*, and assigned it to cube ***r*** as

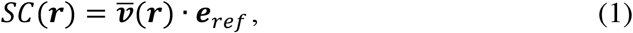

which takes a value of −1 to 1. Green and orange arrows in Figure 9A are vectors 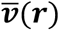 that satisfy the following two conditions:

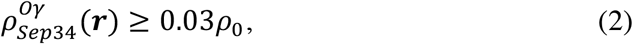

and

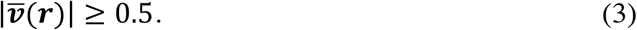

**Figure 9.**
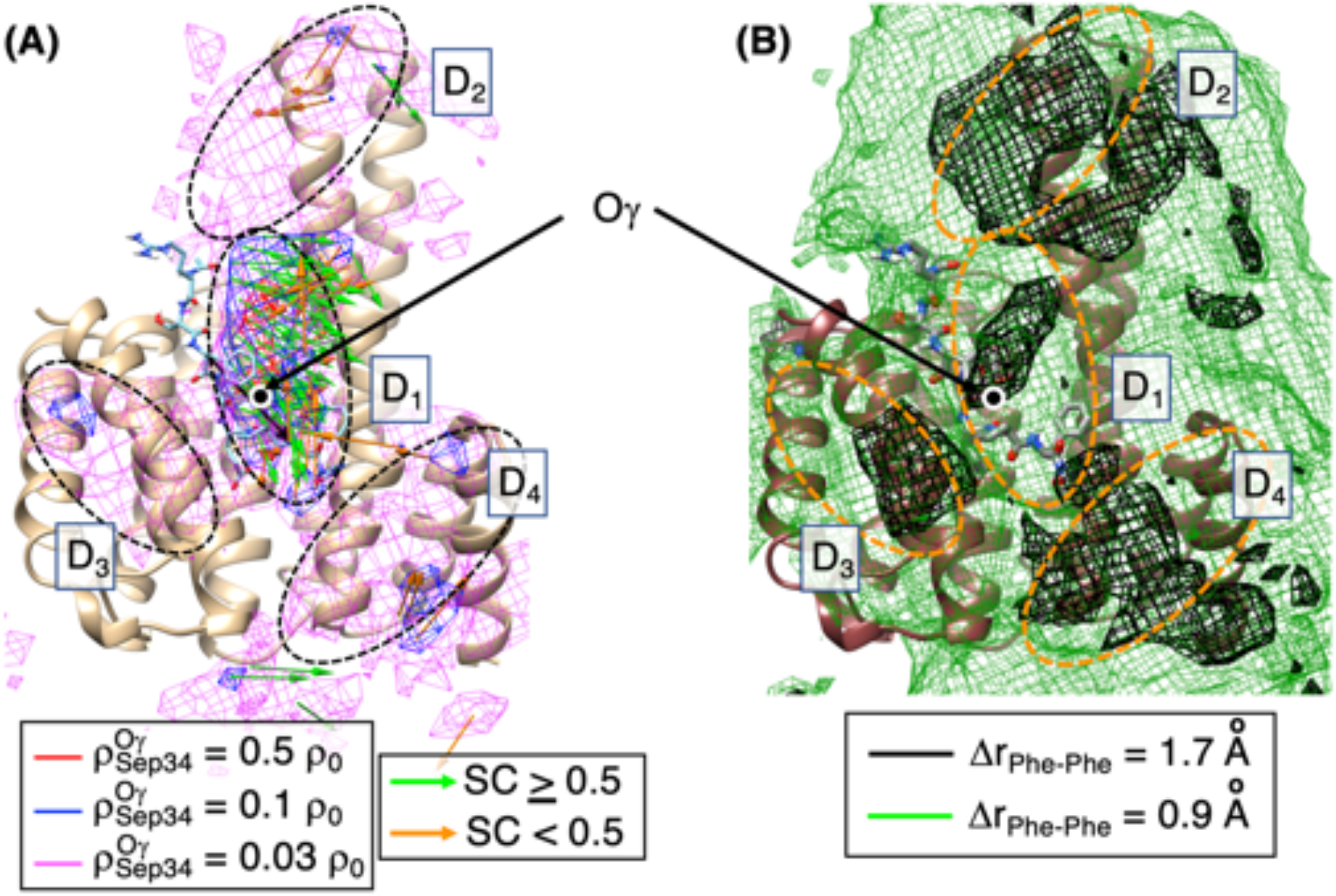
In both panels, the structure which is shown is crustal structure of pMLF1/14-3-3ε, and positions of clusters *D*_1_,.., *D*_4_, which were introduced in Figure 5A, are shown as broken-line circles. Positions of O*γ* atom of Sep 34 of pMLF1 are shown by filled circles with label “O*γ*”. (A) Averaged vectors 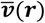 in cubes ***r*** satisfy Eqs. 2 and 3. Red-, blue-, and magenta-colored contours are the same as those in Figure 5A, although 14-3-3ε is viewed in different orientation from Figure 5 to show the positiondependency of 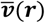. Green and orange arrows respectively represent vectors with *SC*(***r***) ≥ 0.5 and *SC*(***r***) < 0.5. Vectors are multiplied by 10.0 to show the vectors clearly: arrow = 10.0 × 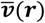. Black arrow points from the Cα atom of Phe 33 to the Cα atom of Phe 37, which shows orientation of pMLF1 in the crystal structure, ***e**_ref_* (B) Spatial patterns of Δ*r*_*Phe–Phe*_(***r***). Black contours show regions of Δ*r*_*Phe–Phe*_(***r***) = 1.7 Å. Green ones show regions of 0.9 Å.

Equation 2 is used to pick cubes involved in high-density regions (clusters *D*_1_,…, *D*_4_ in Figure 5). Also, Eq. 3 is used to display well ordered 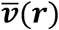. Apparently, only cluster *D*_1_ exhibited remarkable orientational ordering of pMLF1. Its majority was occupied by *SC*(***r***) ≥ 0.5 (green arrows). In contrast, cubes with *SC*(***r***) < 0.5 were rare and sparse. We infer that the orientation of pMLF1 was ordered only in cluster *D*_1_: In other words, the pMLF1 orientation is ordered as reaching the high-density region (i.e., low free-energy region) to be beneficial for binding to 14-3-3ε.

Next, we calculated a ligand-conformational variety, Δ*r*_*Phe–Phe*_(***r***), of pMLF1 at each cube ***r***. Details for Δ*r*_*Phe–Phe*_(***r***) are presented in section 9 of SI. Larger Δ*r*_*Phe–Phe*_(***r***) in a cube ***r*** are associated with a larger the ligand–conformational variety in the cube. We note that Δ*r*_*Phe–Phe*_(***r***) is not linked directly to dynamic chain flexibility of pMLF1 because the conformational ensemble from the GA-guided mD-VcMD is an equilibrated one at 300 K.

Figure 9B demonstrates the spatial patterns of Δ*r*_*Phe–Phe*_(***r***) at two contour levels: Δ*r*_*Phe–Phe*_ = 0.9 Å and Δ*r*_*Phe–Phe*_ = 1.7 Å (green and black contours, respectively). Apparently, 14-3-3ε was surrounded widely by the featureless contours of Δ*r*_*Phe–Phe*_ = 0.9 Å. By contrast, the regions of Δ*r*_*Phe–Phe*_ = 1.7 Å were localized. Figure S12 presents the regions of Δ*r*_*Phe–Phe*_ = 1.7 Å as well as the density maps of 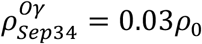 from two orientations. Apparently, clusters *D*_2_, *D*_3_ and *D*_4_ overlapped the regions of Δ*r*_*Phe–Phe*_ = 1.7 Å well. It is likely that a cube occupied by randomized pMLF1 conformations is characterized by less orientation ordering and a wide ligand– conformational variety.

However, the situation of cluster *D*_1_ differs from those of *D*_2_, *D*_3_ and *D*_4_. As shown in Figure 9A, cluster *D*_1_ was characterized by a well-ordered orientation. Figure 11B demonstrates that the contours of Δ*r*_*Phe–Phe*_ = 1.7 Å overlapped with only a part of cluster *D*_1_, and that the high-density spots *E*_1_ and *E*_2_, which were characterized by the higher contour level 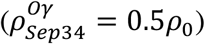 in Figure 6A, were not involved in the Δ*r*_*Phe–Phe*_ = 1.7 Å contours. The O*γ* atom of Sep 34 in the crystal structure is involved in spot *E*_1_. Therefore, we concluded that cluster *D*_1_, to which the well-ordered ligand orientations were assigned, is not characterized by the large ligand-conformational variety.

### Site-dependence of residue interactions between pMLF1 and 14-3-3ε

The four distances *r*^*μ*^ (*μ* = *Phe*33, *Sep*34, *Glu*35, *Phe*37) were defined for each snapshot for Figure 4. Investigating spatial patterns of these distances around 14-3-3ε is interesting. For this purpose, we calculated thermal average 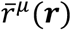 of these distances for each cube ***r*** for the pMLF1/14-3-3ε system according to the procedure explained in section 10 of SI. The thermal average 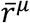 assigned to cube ***r*** is computed using snapshots for which the O*γ* atom of Sep 34 is detected in cube ***r***.

Figure 10A displays the overall spatial patterns of 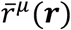 around 14-3-3ε. The red contours show regions with 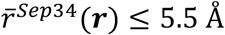. One might consider that the 5.5 Å value is larger than the limit distance for a salt bridge. However, the contours are assigned to the O*γ* atom of Sep 34, not to three O atoms of the phosphate group. Therefore, we increased the limit distance. We also set 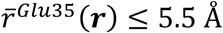 as described before. The limits for Phe 33 and Phe 37 of pMLF1 were set to 7 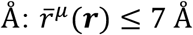, which is larger than the value for a hydrophobic contact because *r*^*γ*^(*i*) was calculated using the ring center (not individual atoms of the ring) of Phe 33 and Phe 37.

**Figure 10.**
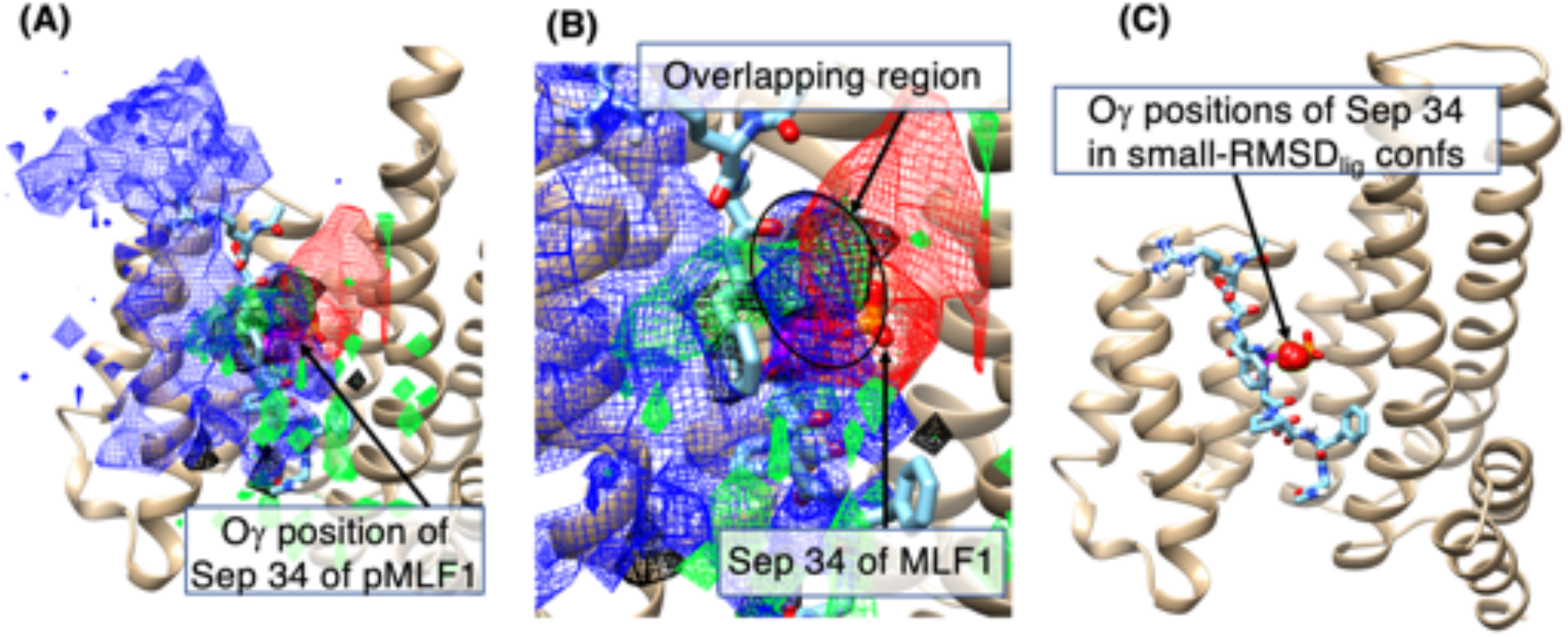
(A) Regions that satisfy 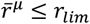. Red-colored contour regions are those for *μ* = *Sep*34 of pMLF1 and *r_lim_* = 5.5 Å. Blue-colored contour regions are those for *μ* = *Phe*33 of pMLF1 and *r_lim_* = 7.0 Å. Black-colored contour regions are those for *μ* = *Glu*35 of pMLF1 and *r_lim_* = 5.5 Å. Green-colored contour regions are those for *μ* = *Phe*37 of pMLF1 and *r_lim_* = 7.0 Å. Black arrow with words “O*γ* position of Sep 34 of pMLF1” denotes its position in crystal structure. (B) Closeup around Sep 34 of pMLF1 in the crystal structure. Four contours overlap around the black-line circle. (C) The O*γ* atomic positions (red spheres show “O*γ* positions of Sep 34 in small-*RMSD_lig_* confs”) of snapshots with *RMSD_lig_* ≤ 1.0 Å. The molecular structure shown is the crustal structure.

Contours for Phe 33 of pMLF1 (blue contour) occupied large regions in Figure 10A, which indicates that Phe 33 can form hydrophobic contacts to Leu 223 and Leu 230 of 14-3-3ε readily. When the O*γ* atom of Sep 34 of pMLF1 was inside of the blue contour regions, Phe 33 was able to form the hydrophobic contact to its interaction partners of 14-3-3ε. This result raises the possibility that these hydrophobic contacts might act as attractive interactions between pMLF1 and 14-3-3ε. Although we have no definite answer to this argument, results of our earlier work examining mD-VcMD^39^ suggest that a long N-terminal tail of a protein might capture its ligand before the ligand binds to the binding site of the receptor (fly casting mechanism).^54–57^ Consequently, the hydrophobic contacts with respect to Phe 33 of pMLF1 might play a role in engendering the most stable complex structure.

Figure 10A shows a large region (red contours) for Sep 34 of pMLF1. This region well overlaps high-density spots *E*_1_ and *E*_2_ (Figure 6). This result is natural because Sep 34 can form salt bridges when Sep 34 approaches the binding partners.

In contrast, the green contours for Phe 37 of pMLF1 were sparse on the surface of 14-3-3ε, which suggests that the hydrophobic interaction of Phe 37 to 14-3-3ε is weaker than that of Phe 33 (blue contours) to stabilize the complex structure. The black contours for Glu 35 of pMLF1 were localized. Therefore, Glu 35 formed the salt bridge only when Glu 35 reached the binding partner Lys 50 of 14-3-3ε.

Figure 10B portrays a closeup image of Figure 10A, particularly addressing Sep 34 of pMLF1 in the crystal structure. Apparently, the four contours overlapped in a narrow region around the Sep 34 position of the crystal structure, as shown in the figure by a black-line circle with the words “Overlapping region”. We emphasize that this narrow region involves the O*γ* atomic position of Sep 34 in the crystal structure. Consequently, the simultaneous formation of the four important intermolecular interactions related to Phe 33, Sep 34, Glu 35, and Phe 37 of pMLF1 are possible only when the O*γ* atom is near the crystal structure position. As presented in Figure 8, Glu 35 shifted somewhat from the crystal-structure position towards the solvent. It is particularly interesting that the black-contours related to Glu 35 in Figures 10A and 10B are the consequence from the shift of Glu 35. Therefore, it is likely that pMLF conformations with the Glu-35 shift belong to the crystal-structure-like complex.

To check the above conjecture further, we selected snapshots with *RMSD_lig_* ≤ 1.0 Å, and displayed the O*γ*-atomic positions of these small-*RMSD_lig_* snapshots in 3D real space. In fact, Figure 10C demonstrates that the O*γ*-atomic positions of those snapshots overlapped well to the crystal-structure position. This result also supports that simultaneous satisfaction of the four intermolecular interactions engenders the crystal-structure-like complex form.

## DISCUSSION

The current study presents a binding process of pMLF1 to 14-3-3ε (Figure 11). Before binding to 14-3-3, pMLF1 exists everywhere around 14-3-3ε at a low concentration (*ρ*_*Sep*34_ = 0.004 *ρ*_0_; dark green contours). It is presumable from the magenta-colored contours (*ρ*_*Sep*34_ = 0.03 *ρ*_0_) that pMLF1 can contact to four regions of the 14-3-3ε surface, *D*_1_,…, *D*_4_, more stably or more frequently than to the other regions. An encounter complex is thereby formed. If pMLF1 binds to *D*_1_ first, then pMLF1 might form a crystal-structure-like complex quickly by adjustment of the conformation.

**Figure 11.**
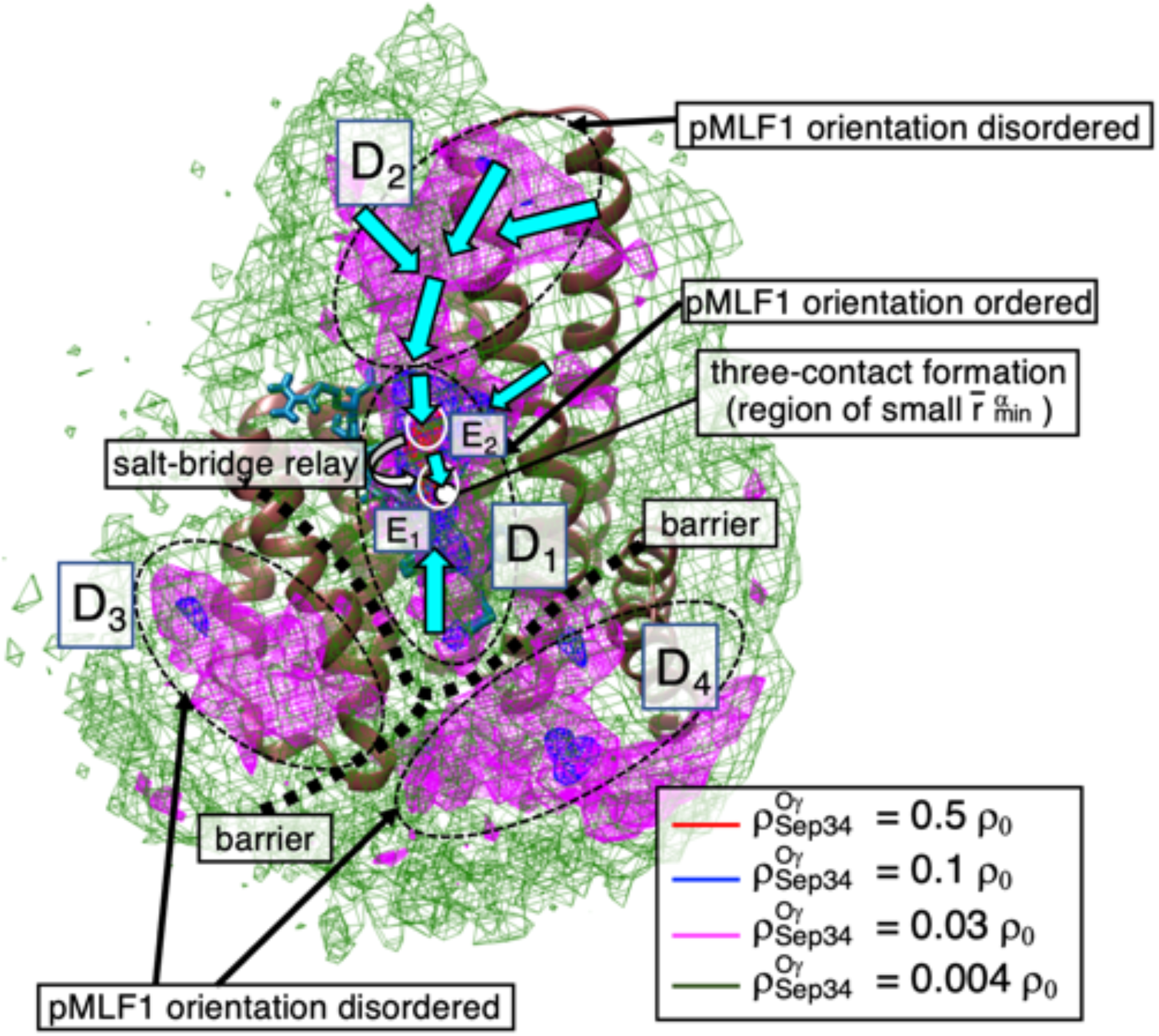
Scheme of binding process of pMLF1 to 14-3-3ε. Contour maps represent *ρ*_*Sep*34_(***r***) at four density levels as for Figure 5. Clusters *D*_1_,…, *D*_4_ are also from Figure 5. Free-energy barriers are shown by thick broken lines. Cyan-colored arrows represent conformational motions of pMLF1 from cluster *D*_2_ to *D*_1_. Molecular orientation of pMLF1 is ordered well in *D*_1_, although it is less ordered in *D*_2_, *D*_3_, and *D*_4_. The position where salt-bridge relay takes place is shown by a curved arrow. White open circles are high-density spots *E*_1_ and *E*_2_. White filled circles show positions of four intermolecular contacts, quantified by 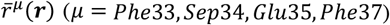, are formed simultaneously.

Because the sum of the regions *D*_2_ + *D*_3_ + *D*_4_ is greater than *D*_1_, it is likely that pMLF1 binds more frequently to *D*_2_, *D*_3_, or *D*_4_ than to *D*_1_. Free-energy barriers exist among three clusters *D*_1_, *D*_3_, and *D*_4_. Therefore, if pMLF1 contacts to *D*_3_ or *D*_4_ first, then the complex should jump over the free-energy barriers to reach *D*_1_. This motion might take a long time. Otherwise, the encounter complex generated in *D*_3_ or *D*_4_ might be dissociated sooner or later.

A scenario for the complex formation presumed from the current study is the following: The residue Phe 33 of pMLF1 might contact the hydrophobic contact partner of 14-3-3ε even when pMLF1 is separate from the binding site of 14-3-3ε (Figure 10A). This contact might act as an attractive interaction between pMLF1 and 14-3-3ε. Then, if the encounter complex is formed in *D_2_* first, then no energy jump is necessary to move to *D*_1_ because the two clusters are connected smoothly without a free-energy barrier. Remember that pMLF1 has no orientational tendency in *D*_2_. When pMLF1 moves to *D*_1_, Sep 34 of pMLF1 meets spot *E*_2_ and binds to Arg 61 and/or Arg 57 of 14-3-3ε. Simultaneously, the molecular orientation of pMLF1 tends to be aligned as in the crystal structure. At that stage, neither the hydrophobic contacts related to Phe 37 of pMLF1 nor the salt bridge related to Glu 35 of pMLF1 might be formed. When pMLF1 goes through the corridor from spot *E*_2_ to *E*_1_, the salt-bridge relay takes place: The salt bridge between Sep 34 of pMLF1 and Arg 61 of 14-3-3ε is broken. Also, that between Sep 34 of pMLF1 and Arg 130 of 14-3-3ε is formed. When Sep 34 fluctuates in spot *E*_1_, which involves the crystal-structure position (white-circle position in Figure 11), Phe 37 forms hydrophobic contacts to 14-3-3ε. Simultaneously, Glu 35 forms the salt bridge to Lys 50, although the salt bridge to Lys 123 might not be formed.

## CONCLUSIONS

The equilibrated distribution functions derived from the GA-guided mD-VcMD suggest a binding process of pMLF1 to 14-3-3ε (Figure 11). Furthermore, the distribution provided interesting aspects such as free-energy barriers among *D*_1_, *D*_3_, and *D*_4_. It also provided pMLF1 orientation ordering and the small ligand–conformational variety in the high-density region and a salt-bridge relay to reach the complex structure similar to the crystal structure. The narrow spot is fundamentally important for crystal structure formation. These aspects are derived naturally from the current simulation without a gap in logic. They are useful to advance other experimental and computational studies.

In our simulation, Glu 35 shifted somewhat toward solvent. It exhibited large positional fluctuations, even in the high-density spot (Figure 8). These positional shift and large fluctuations were induced by absence of the salt bridge to Lys 123 of 14-3-3ε, which is formed in the crystal structure (PDB ID: 3ual). Our simulation study supports experimental data showing that mutation E35A of pMLF1 does not alter *K_D_* for the pMLF1/14-3-3ε complex.^7^ Results of our simulation also suggest that large fluctuations of Glu 35 affect the hydrophobic interaction of Phe 37 with the sidechain stem of Lys 50. Mutagenesis K50A of 14-3-3ε might provide insightful information supporting our inference.

Results of the current study demonstrated that the GA-guided mD-VcMD can treat a very flexible ligand molecule without restraining the molecular flexibility. It is noteworthy that this method can search a wide conformational space in which the ligand covers not only the vicinity of the binding site of the receptor, but also regions distant from the binding site. That is true because the RCs were set to search the wide 3D real space. Although the best selection of the reaction coordinates is not known a priori, the overlap of basins in the RC space is, in general, resolved by increasing the space dimensionality. For this reason, the introduction of multiple RCs was useful in this study to sample the highly flexible chain, pMLF1, binding to the receptor. In contrast, incrementation of the space dimensionality makes sampling data sparse in multi-dimensional RC space. This study has demonstrated that 3D is appropriate as a tradeoff between the basin-overlap resolution and sparse sampling.

The molecular binding scheme for an intrinsically disordered region/protein (IDR/IDP) has variety: Coupled folding and binding,^56,58^ fuzzy complexes,^59^ multimodal complexes,^53^ and dynamic complexes.^60^ The current study demonstrated that pMLF1 exists in two high-density spots *E*_1_ and *E*_2_ (Figure 6A). These multiple-state features might also be related to the property of IDR/IDP.

Recently, to investigate intermolecular interactions, we performed multicanonical molecular dynamics (McMD) of an intrinsically disordered peptide, p53 C-terminal domain (p53CTD), binding to its receptor protein (S100B).^37^ It is particularly interesting that results of this study suggest that the degree of disorder of p53CTD increases when binding to S100B. We designated this emerging state upon binding an *extra–disorder* state. For this study, we analyzed whether the extra-disorder state existed or not by comparing the ligand conformational variety between the bound and unbound states. Nevertheless, we were unable to clarify any clue to the extra-disordered state because the upper limits of 40 Å for RCs (Table S2) were still too short to sample pMLF1 conformations entirely free from 14-3-3ε: that is, a pMLF1 snapshot that was not touching 14-3-3ε can achieve touching by rotating pMLF1 (data not shown). Therefore, sampling of the unfolded state was insufficient. As a subject of future study, we set the RC limits to values that are sufficiently large to generate a ligand completely free from the receptor. Furthermore, it is noteworthy that this increment of RCs leads the GA-guided mD-VcMD to calculate *K_D_* directly by calculating the probabilities for both the bound and free states.

The density contour map obtained from the current study can be expected to produce many insights. For instance, a molecular modification that changes the corridor width between spot *E*_1_ and *E*_2_ alters the kinetic constants. It therefore affects the complex association and dissociation constants. The current simulation also suggests a reason why the mutation E35A did not alter *K_D_*. These insights might pave the way to new approaches for drug design. We emphasize that the large-scale conformational ensemble equilibrated at a room temperature, which covers the free, encounter-complex, and the most-stable states, is useful to widen the research area for drug discovery.

## ACKNOWLEDGEMENTS

The authors thank Prof. Norio Ozaki in Nagoya University Graduate School of Medicine for his valuable discussion of the functional roles of the 14-3-3 proteins which interact with the phosphorylated sequences. J. H. was supported by JSPS KAKENHI Grant No. 16K05517 and by the Development of Core Technologies for Innovative Drug Development based upon IT from Japan Agency for Medical Research and Development, AMED. J. H. and K. K. were supported by the HPCI System Research Project (Project IDs: hp190017, hp190018, hp190027, hp200025, hp200063, and hp200090). The mD-VcMD was performed on the TSUBAME3.0 supercomputers at the Tokyo Institute of Technology. K. K. was also supported by JSPS KAKENHI Grant No. 16K18526. H. N. was supported by a Grant-in-Aid for Scientific Research on Innovative Areas (24118008) and a Grant-in-Aid for Challenging Exploratory Research (16K14711) from JSPS. N. K. was supported by JSPS KAKENHI Grant (Number JP20H03229). It was performed in part under the Cooperative Research Program of the Institute for Protein Research, Osaka University, CR-19-05 and CR-20-05.

## Supporting Information

**Figure S1.**
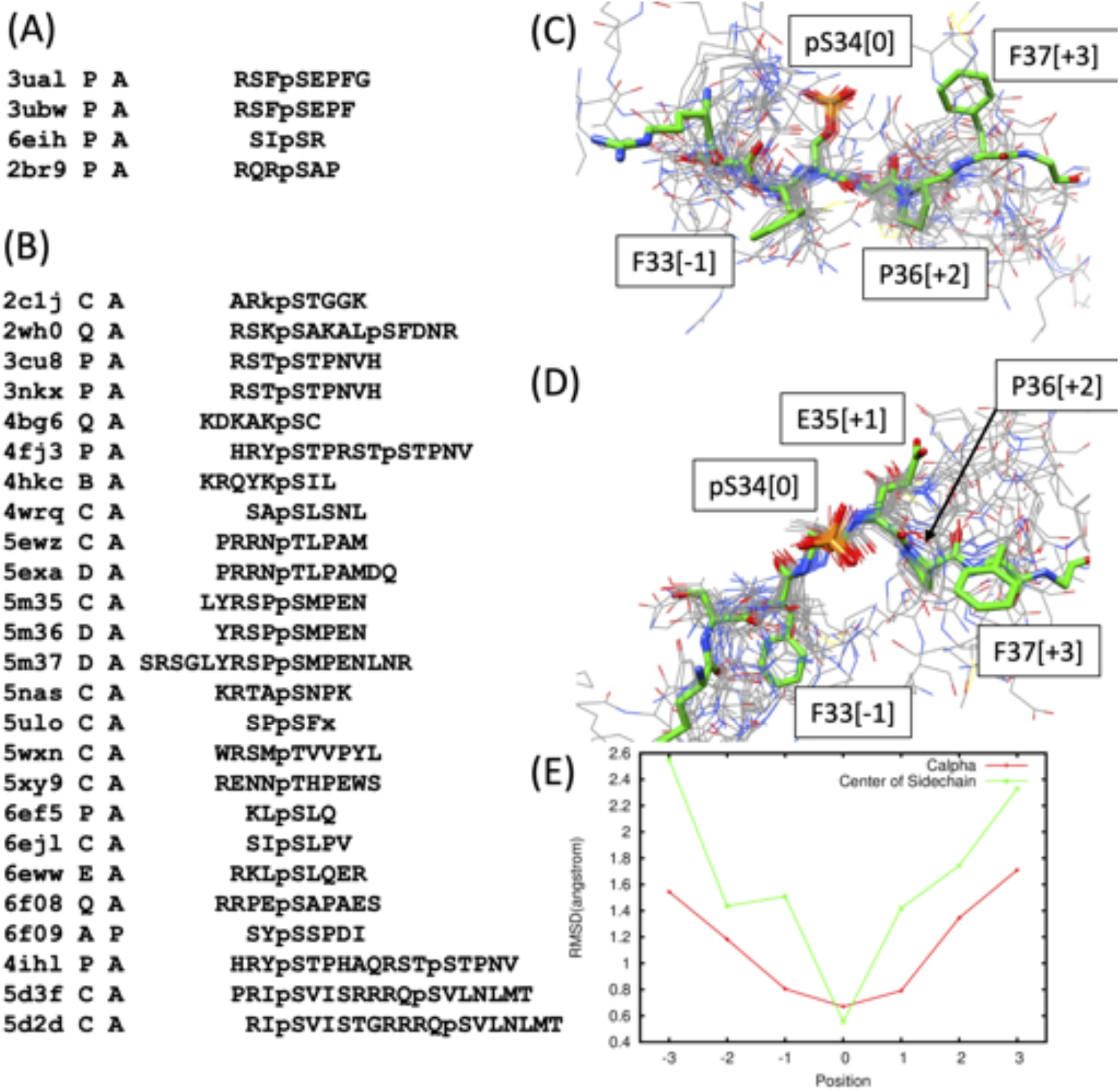
(A) Four structurally known phosphorylated peptides bound to human 14-3-3*ε* protein. PDB IDs, chain IDs of binding peptide, chain IDs of 14-3-3 protein and amino acid sequences of binding peptides are shown. pS and pT are phosphorylated serine and threonine residues, respectively. (B) 25 structurally known phosphorylated peptides bound to human 14-3-3*ζ* protein. Sequence identity between 14-3-3*ε* and 14-3-3*ζ* is 67 %. (C) A view of the superimposed 3D structures of binding peptides to 14-3-3*ε* or 14-3-3*ζ* proteins. They were superimposed by their bound 3D structures of 14-3-3 proteins on A chain (14-3-3*ε*) of PDB ID:3ual, using the program MATRAS.^[1]^ The peptide structure of P chain (i.e., pMLF1) of PDB ID:3ual is shown in green. The residue numbering accords to UniProt (M.MLF1_HUMAN; https://www.uniprot.org/uniprot/P58340). Numbers in square brackets are residue ordinal numbers counted from the phosphorylated residue. (D) A view of the superimposed 3D structures from a different orientation. (E) RMSD plots of the superimposed 28 peptides with the P chain of PDB ID:3ual. Red line shows RMSDs of the corresponding C*α* atoms, and the green line those of center of sidechain atoms.

**Figure S2.**
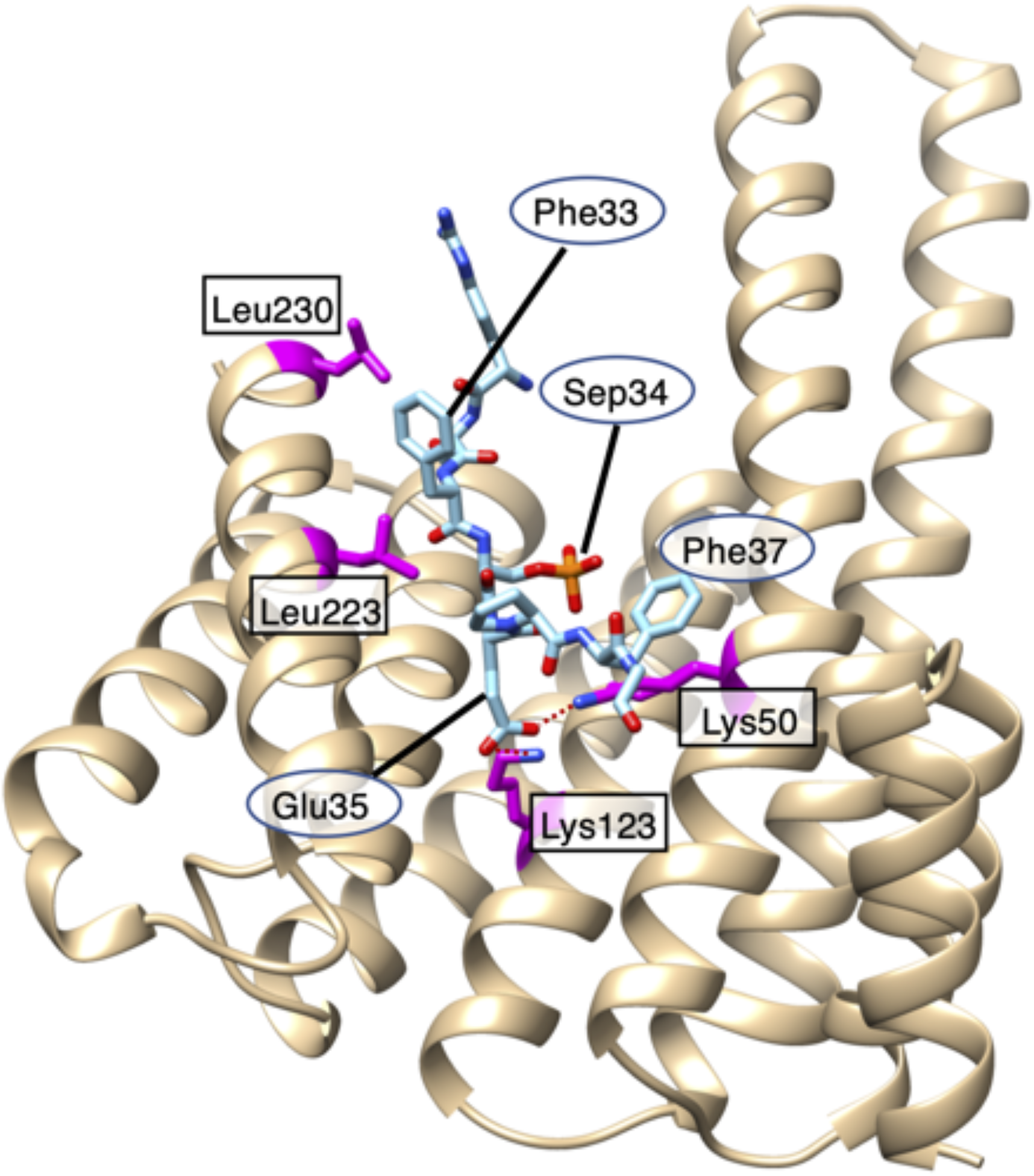
14-3-3*ε*/MLF1 complex structure solved by X-ray crystallography, where MLF1 and 14-3-3*ε* are shown in cyan and ocher, respectively. Residues Phe 33, Sep 34, Glu 35, and Phe 37 belong to MLF1, and Lys 50, Lys 123, Leu 223, and Leu 230, shown by magenta, do to 14-3-3*ε*. Salt bridges from Glu 35 of MLF1 to Lys 50 and Lys 123 of 14-3-3*ε* are shown by brown-colored dotted lines. Two hetero sidechains are shown for Lys 50.

### Section 1. Basic method to define a reaction coordinate (RC)

Consider two atom groups 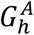 and 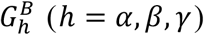 in a molecular system. The RC, *λ*^(*h*)^, is defined by the distance between centers of mass of 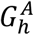 and 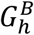 (Fig. S3). Superscripts *A* and *B* indicate simply that two atom groups are pairing to define *λ*^(*h*)^, and then, one can exchange the superscripts as: 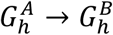 and 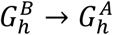 without changing the value of *λ*^(*h*)^.

**Figure S3.**
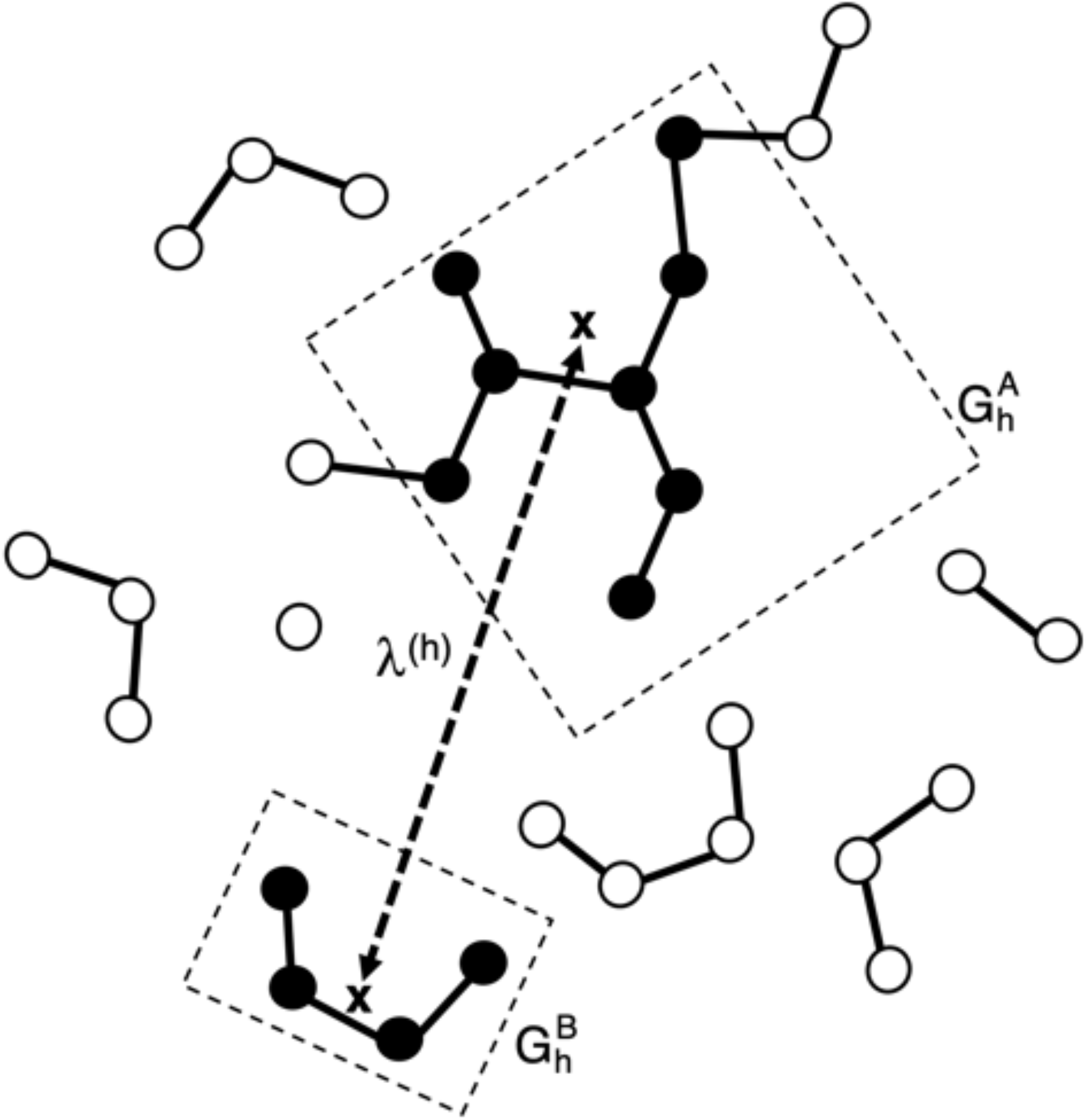
Two atom groups, 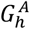 and 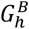, are indicated by two rectangles, where in-group atoms are shown by black filled circles. Center of mass of each atom group is presented by a cross. The distance between the two centers of mass is *λ*^(*h*)^ (broken-line with arrows).

### Section 2. Generation of pMLF1/14-3-3*ε* and npMLF1/14-3-3*ε* systems

To generate the pMLF1/14-3-3*ε* system, first, the X-ray crystal structure of the complex of 14-3-3*ε* with pMLF1 (i.e., MLF1 phosphorylated at the 34-th serine residue; see the main text for the amino-acid sequence), was taken from the PDB data (ID: 3ual).^2^ Residue ordinal numbers used in this study accord to those in the paper,^2^ which are not the same as those used in the PDB data. The receptor 14-3-3*ε* forms a homodimer in the unbinding state, as shown in Fig. S4A schematically. The ligand pMLF1 binds to each of the two 14-3-3*ε* molecules (Fig. S4B), where Sep 34 (the 34th serine residue phosphorylated) of pMLF1 forms salt bridges to Arg 57 and Arg 130 of 14-3-3*ε* in the crystal structure (Fig. S2). To reduce the computational load, in the present study, we used a single complex of 14-3-3*ε* with pMLF1 (Fig. S4C). The N- and C-termini of both 14-3-3*ε* and pMLF1 were capped by acetyl and N-methyl groups, respectively. The single complex taken from 3ual was immersed in a periodic water box (86.2 × 86.2 × 90.8 Å^3^) under periodic boundary condition, and sodium and chlorine ions were introduced to neutralize the net charge of the entire system at a physiological ionic concentration. The resultant system consists of 65,776 atoms (3,700 atoms for 14-3-3*ε*; 137 atoms for pMLF1; 20,612 molecules for water; 46 chlorine ions and 57 for sodium ions).

**Figure S4.**
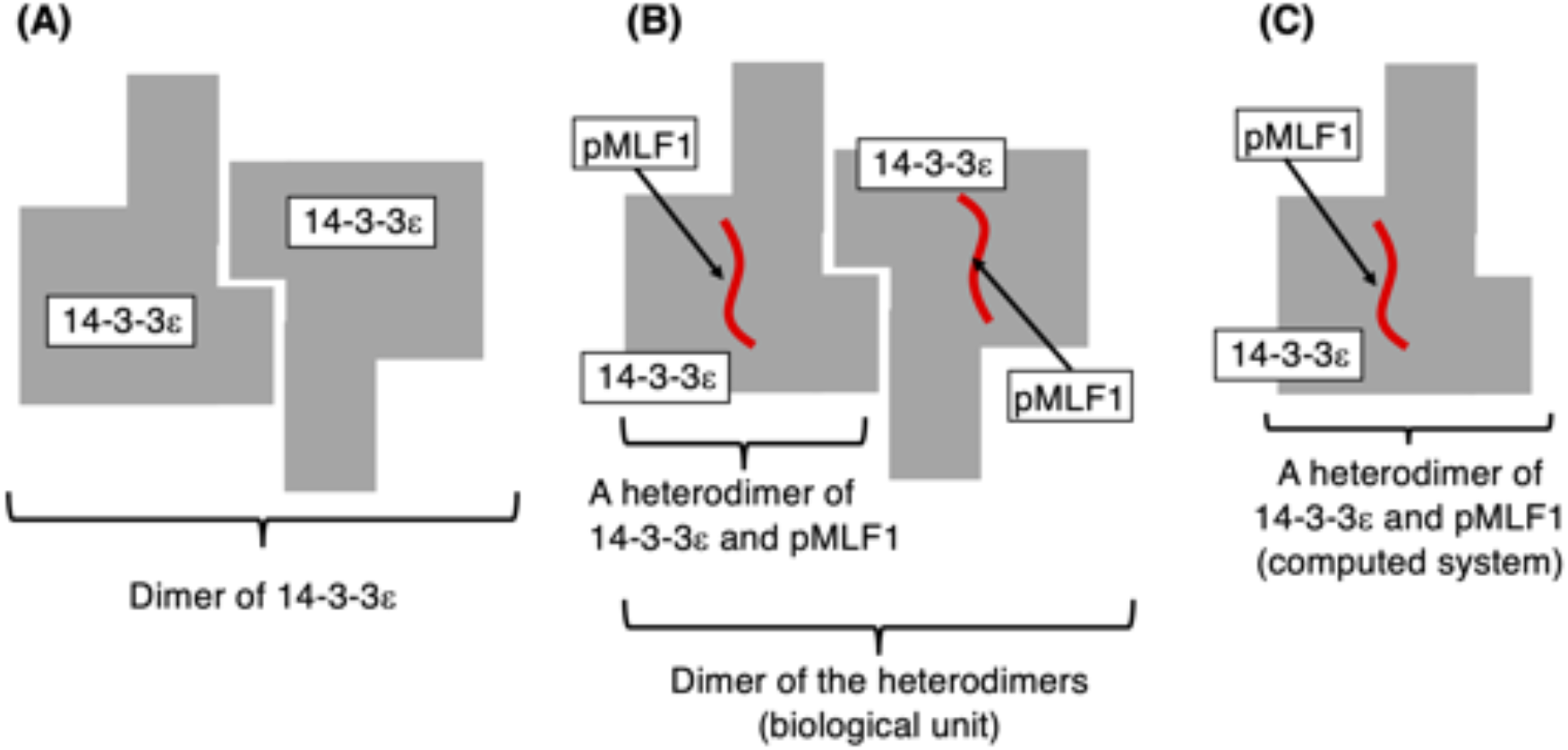
Shaded polygon and red-colored string are receptor 14-3-3*ε* and ligand pMLF1, respectively. (A) 14-3-3*ε* dimer in the unbound state. (B) Dimer of heterodimers [pMLF1/14-3-3 *ε*]_2_ formed when pMLF1 binds to 14-3-3*ε*. (C) Computed system.

Although pMLF1 used for the experiment^2^ was a segment of 14 residues long, the two N- and four C-terminal residues were not determined in the experiment. Then, we treated the eight-residue determined segment in the present study. The receptor 14-3-3*ε* consists of 230 amino-acid residues (see PDB: 3ual). We refer to the current system consisting of the capped pMLF1 and 14-3-3*ε* as the pMLF1/14-3-3*ε* system, whether the two molecules form a complex or not during simulation.

We also computed a system of 14-3-3*ε* and npMLF1 (the non-phosphorylated MLF1; see the main text for the amino-acid sequence), and we refer to this system as an npMLF1/14-3-3*ε* system. This system was generated by replacing Sep 34 of pMLF1 in the pMLF1/14-3-3*ε* system by serine. Then, the numbers of ions and water molecules were modified to neutralize the net charge of the system. The resultant system consists of 65,771 atoms (3,700 atoms for 14-3-3*ε*; 134 atoms for npMLF1; 20,612 molecules for water; 46 chlorine ions and 55 for sodium ions).

The pMLF1/14-3-3*ε* complex structure generated above is the same as the crystal structure. On the other hand, a stable complex is not reported for the npMLF1/14-3-3*ε* system. However, the npMLF1/14-3-3 *ε* complex structure generated above adopts the crystal structure of the pMLF1/14-3-3*ε* complex except for the sidechain of Ser 34. Thus, for convenience, this structure is also called the crystal structure in this study.

For each of the pMLF1/14-3-3 *ε* and npMLF1/14-3-3 *ε* systems, after a short energy minimization of the complex structure, a short constant-volume and constant-temperature (300 K) simulation (NVT simulation) was performed. Then, the system size was relaxed by a constant-pressure (1 atm) and constant-temperature simulation (NPT simulation). The resultant box size was 84.70 × 84.67 × 89.24 Å^3^ and 84.72 × 84.69 × 89.27 Å^3^ for the pMLF1/14-3-3 *ε* and npMLF1/14-3-3*ε* systems, respectively.

We aim to search the conformational space of both systems starting from randomized ligand’s conformations of pMLF1 or npMLF1. However, the obtained structures from the NPT simulation were still close to the crystal structure (data not shown). Thus, we generated the randomized conformations as explained in section 3 of SI below. The word “initial conformation” used in the present study represents the randomized conformation where pMLF1 or npMLF1 is apart from the pMLF1 binding site of 14-3-3*ε*.

### Section 3. Generation of initial conformations

As shown in Fig. 1 of main text and Table S1, we introduced three RCs, *Λ*^(*α*)^, *λ*^(*β*)^, and *λ*^(*γ*)^. To generate randomized conformations of ligand (pMLF1 or npMLF1), we performed a simulation with applying weak repulsive forces along the three RCs so that the RCs fell in the ranges of 38 Å ≤ *λ*^(*h*)^ ≤ 40 Å (*h* = *α,β,γ*). Then, for both systems, we performed 256 runs starting from the conformation obtained from the NPT simulation with different initial velocities of atoms, and used the final snapshots for the initial conformations of the first iteration of GA-guided mD-VcMD. In the resultant snapshots, pMLF1 or npMLF1 was distant from the binding site of 14-3-3*ε*, which were slightly contacting to or unbound from 14-3-3*ε*. Figure S5 of SI displays some of the initial conformations randomly picked from the 256 initial conformations.

**Figure S5.**
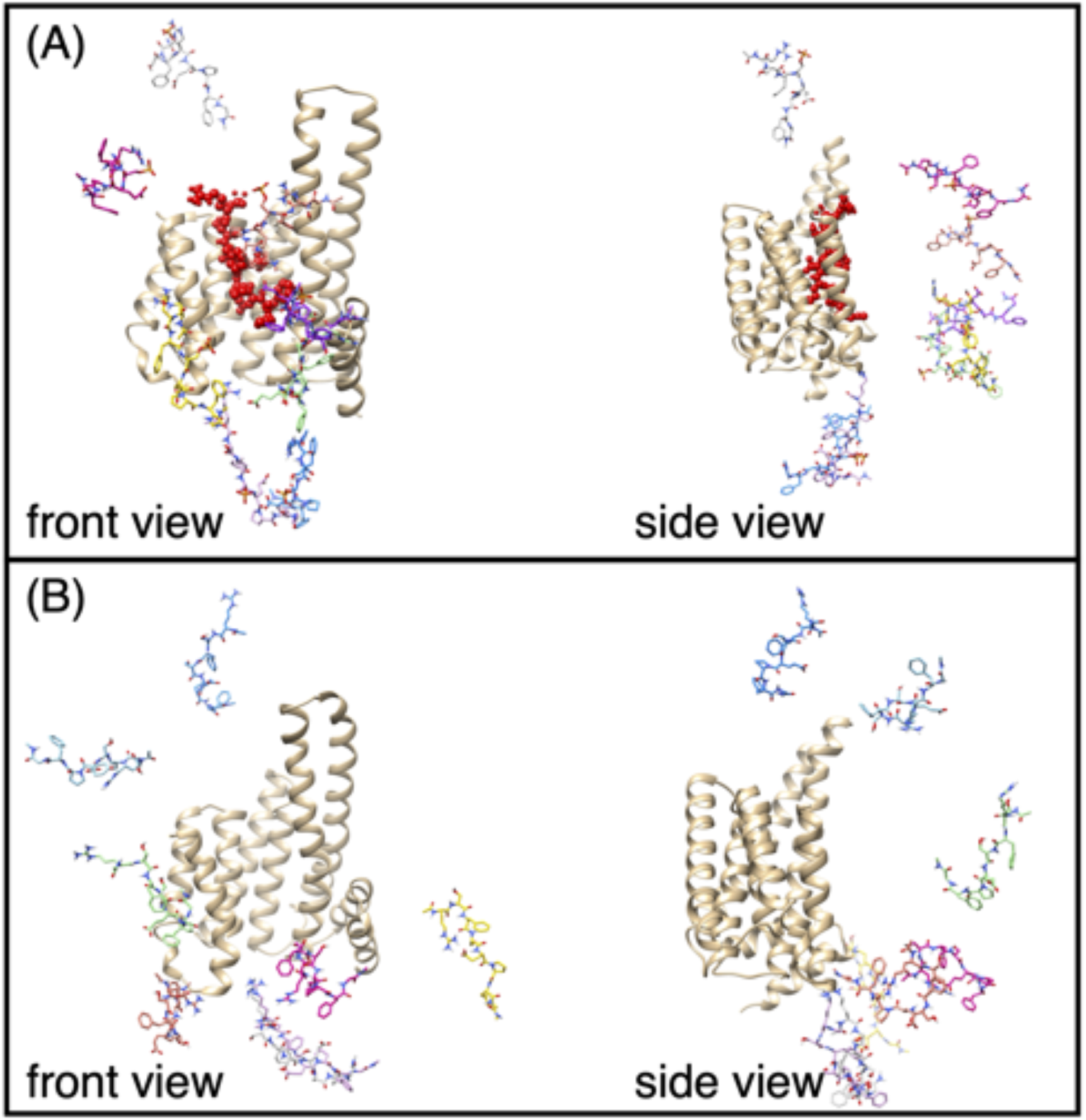
The initial conformations of GA-guided mD-VcMD for (A) pMLF1/14-3-3*ε* and (B) npMLF1/14-3-3*ε* systems. Eight conformations picked randomly from 256 initial ones are shown. Red-colored model in panel (A) is pMLF1 in the crystal structure. Note that this red-colored model is not used for the initial conformation of the GA-guided mD-VcMD. Solvent was omitted in this figure.

### Section 4. Iterative mD-VcMD simulations and parameters used

GA-guided mD-VcMD consists of iterative simulations, through which the conformational ensemble converges on an equilibrated one at a simulation temperature (300 K). We repeated 30 and 14 iterations for the pMLF1/14-3-3*ε* and npMLF1/14-3-3*ε* systems, respectively. To sample carefully the electrostatic interactions between Sep 34 of pMLF1 and charged residues in 14-3-3*ε*, we did more iterations for pMLF1/14-3-3*ε* than npMLF1/14-3-3*ε*. For both systems, a set of 256 runs were bundled in an iteration. Each of 256 runs in an iteration was performed for 2 × 10^6^ steps (4 ns). Thus, the total simulation length was 30.720 μs (= 30 × 256 × 4 ns) and 14.336 μs (= 14 × 256 × 4 ns) for pMLF1/14-3-3*ε* and npMLF1/14-3-3*ε*, respectively. A snapshot was stored every 1 × 10^5^ steps (200 ps), yielding 153,600 and 71,680 snapshots for pMLF1/14-3-3*ε* and npMLF1/14-3-3*ε*, respectively.

Here, we provide actual values of parameters which control the simulations. The atom groups adopted for the current study to define the three RCs are listed in Table S1. Also see Fig. 1 of main text, which graphically explains the atom groups and RCs. Table S2 lists parameters *n_vs_*(*h*), 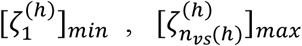, and Δ*λ*^(*h*)^, and Table S3 does actual setting of zones: 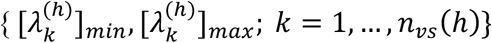. An interzone transition was attempted once every 20 *ps*: The IVT transition interval was set to 1 × 10^4^ steps. See Ref. 3 for meaning of parameters.

**Table S1.** Atom groups to define three RCs.

| RC | Atom Group<sup>a)</sup> |  |
| --- | --- | --- |
| | $G_h^A$ | $G_h^B$ |
| $\lambda^{(\alpha)}$ | Arg 130-Glu 134<sup>b)</sup> | Arg 31 |
| $\lambda^{(\beta)}$ | Leu 173-Val 179 | Sep 34/Ser 34<sup>c)</sup> |
| $\lambda^{(\gamma)}$ | Ala 48-Val 52 | Phe 37 |

<sup>a)</sup> Two atom groups are expressed as $G_h^A$ or $G_h^B$ ( $h = \alpha, \beta, \gamma$ ) to define $\lambda^{(h)}$ : $\lambda^{(h)}$ is defined by the distance between the centers of mass of $G_h^A$ and $G_h^B$ . Superscripts $A$ and $B$ are assigned to 14-3-3ε and pMLF1 (or npMLF1), respectively.
<sup>b)</sup> “ $X$ $n_1$ - $Y$ $n_2$ ” represents all mainchain heavy and $C\beta$ atoms from amino-acid $X$ at the $n_1$ -th residue to amino-acid $Y$ at the $n_2$ -th residue.
<sup>c)</sup> Sep 34 and Ser 34 are the phosphorylated serine of pMLF1 and the serine of npMLF1 at the 34-th residue, respectively.

**Table S2.**
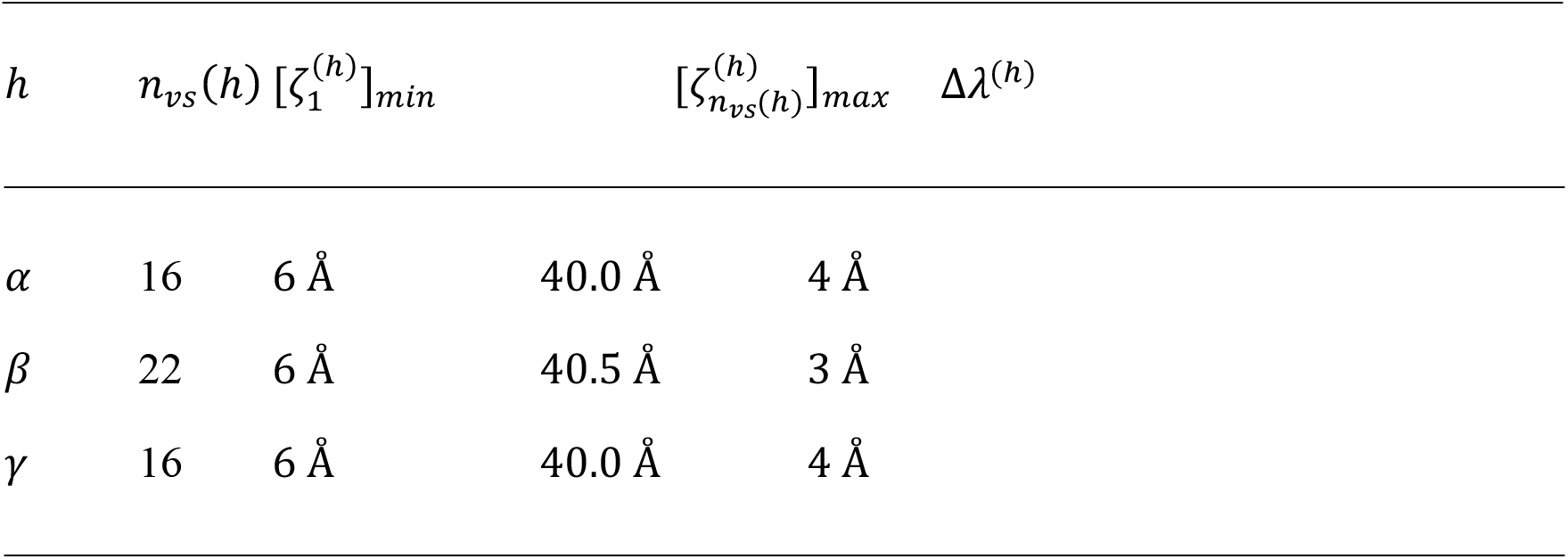
Parameters^a)^ for RC-space division

**Table S3.**
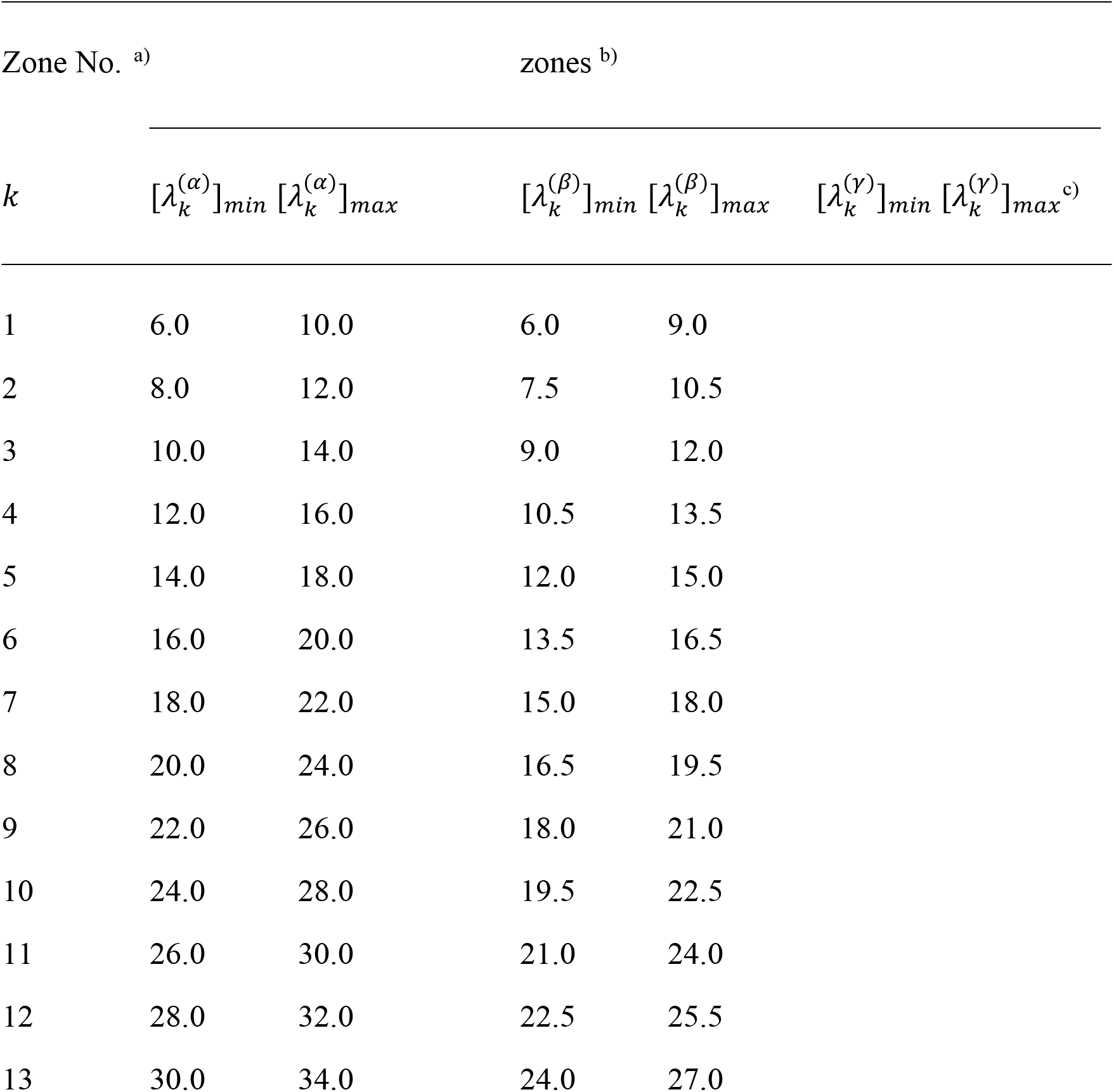
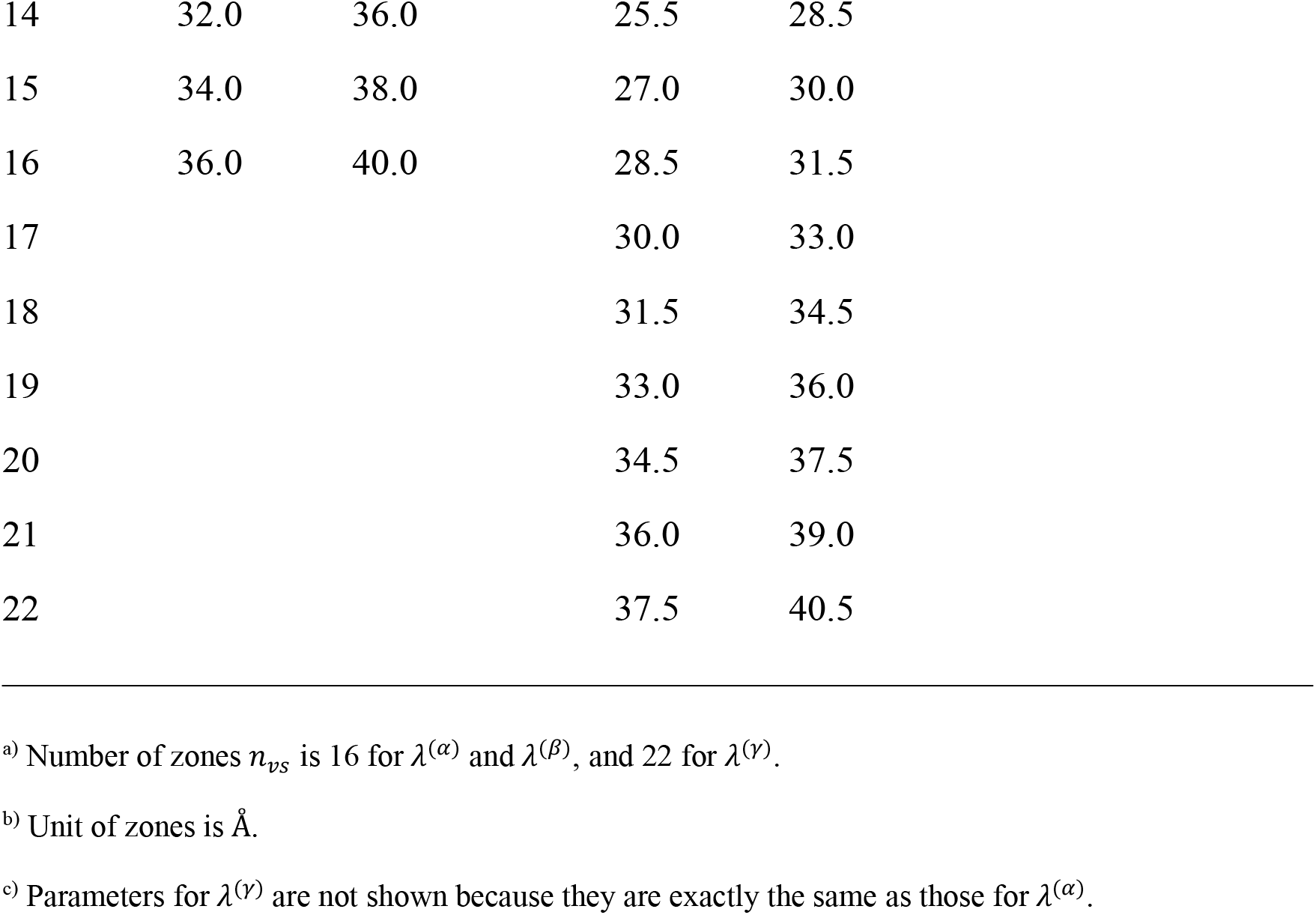
Setting of zones.

| Zone No. <sup>a)</sup> | zones <sup>b)</sup> |  |  |  |  |  |
| --- | --- | --- | --- | --- | --- | --- |
| $k$ | $[\lambda_k^{(\alpha)}]_{min}$ | $[\lambda_k^{(\alpha)}]_{max}$ | $[\lambda_k^{(\beta)}]_{min}$ | $[\lambda_k^{(\beta)}]_{max}$ | $[\lambda_k^{(\gamma)}]_{min}$ | $[\lambda_k^{(\gamma)}]_{max}$ <sup>c)</sup> |
| 1 | 6.0 | 10.0 | 6.0 | 9.0 |  |  |
| 2 | 8.0 | 12.0 | 7.5 | 10.5 |  |  |
| 3 | 10.0 | 14.0 | 9.0 | 12.0 |  |  |
| 4 | 12.0 | 16.0 | 10.5 | 13.5 |  |  |
| 5 | 14.0 | 18.0 | 12.0 | 15.0 |  |  |
| 6 | 16.0 | 20.0 | 13.5 | 16.5 |  |  |
| 7 | 18.0 | 22.0 | 15.0 | 18.0 |  |  |
| 8 | 20.0 | 24.0 | 16.5 | 19.5 |  |  |
| 9 | 22.0 | 26.0 | 18.0 | 21.0 |  |  |
| 10 | 24.0 | 28.0 | 19.5 | 22.5 |  |  |
| 11 | 26.0 | 30.0 | 21.0 | 24.0 |  |  |
| 12 | 28.0 | 32.0 | 22.5 | 25.5 |  |  |
| 13 | 30.0 | 34.0 | 24.0 | 27.0 |  |  |

|  |  |  |  |  |
| --- | --- | --- | --- | --- |
| 14 | 32.0 | 36.0 | 25.5 | 28.5 |
| 15 | 34.0 | 38.0 | 27.0 | 30.0 |
| 16 | 36.0 | 40.0 | 28.5 | 31.5 |
| 17 |  |  | 30.0 | 33.0 |
| 18 |  |  | 31.5 | 34.5 |
| 19 |  |  | 33.0 | 36.0 |
| 20 |  |  | 34.5 | 37.5 |
| 21 |  |  | 36.0 | 39.0 |
| 22 |  |  | 37.5 | 40.5 |

a) Number of zones $n_{vs}$ is 16 for $\lambda^{(\alpha)}$ and $\lambda^{(\beta)}$ , and 22 for $\lambda^{(\gamma)}$ .
b) Unit of zones is Å.
c) Parameters for $\lambda^{(\gamma)}$ are not shown because they are exactly the same as those for $\lambda^{(\alpha)}$ .

**Figure S6.**
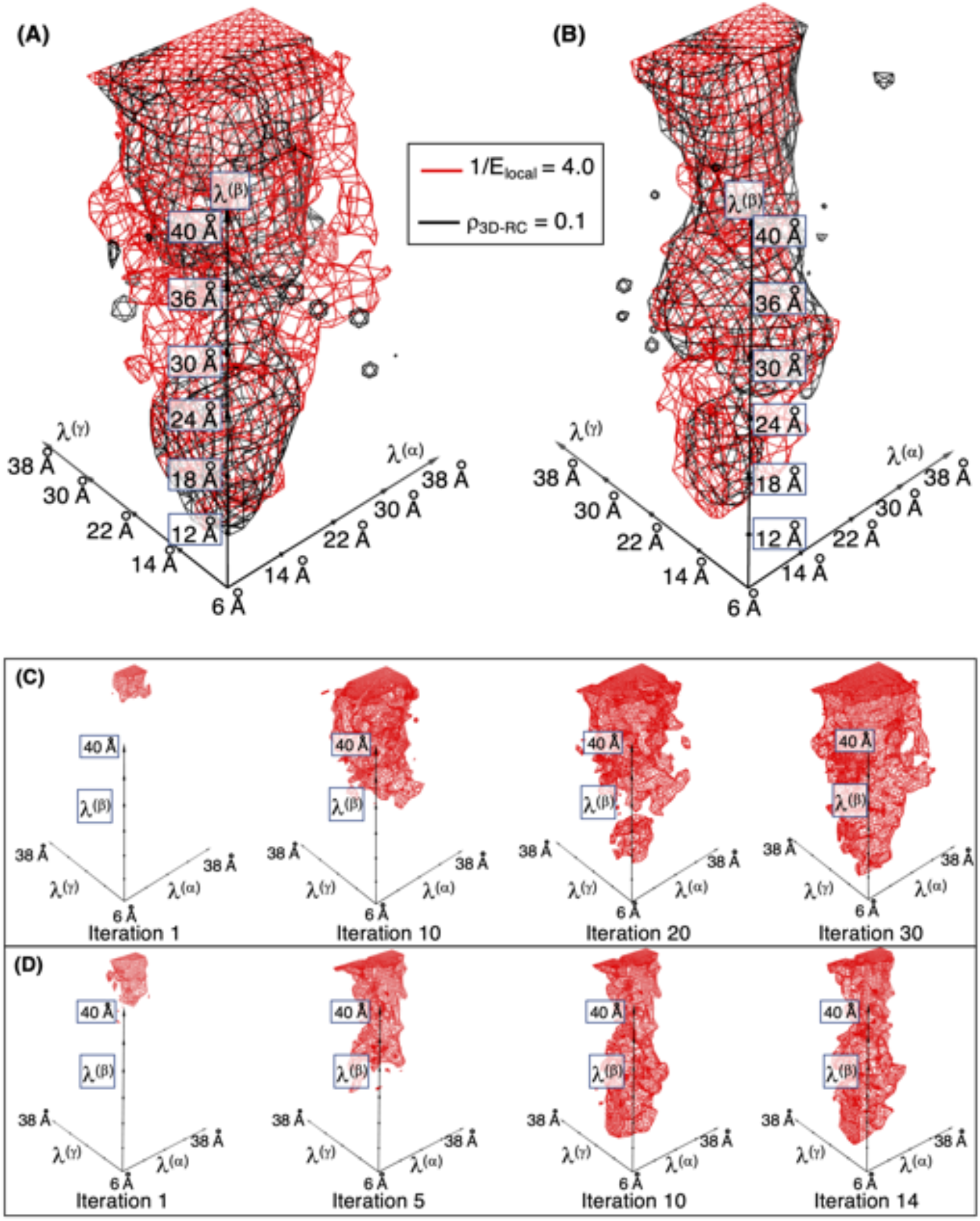
Accuracy of obtained distribution *ρ*_3*D–RC*_(***λ***) assessed by an objective function *E_local_* (***λ***)^-1^ introduced in our previous work.^3^ Benefit of this function is: The accuracy can be accessed locally in the RC space. The larger the *E_local_* (***λ***)^-1^ at a position I, the more accurate the *ρ*_3*D–RC*_(***λ***) at the position. This figure is presented at contour level of *E_local_*(***λ***)^-1^ = 4. Panels (A) and (B) are for pMLF1/14-3-3*ε* and npMLF1/14-3-3*ε*, respectively. We have not yet had an appropriate threshold to assess *E_local_*(***λ***)^-1^ because we have little experience with GA-guided VcMD. However, this figure indicated that regions, to which relatively large *ρ*_3*D–RC*_(***λ***) are assigned (*ρ*_3*D–RC*_ ≥ 0.1), are covered by contours of *E_local_*(***λ***)^-1^ ≥ 4. This is an expected result because the thermodynamically important RC regions should have more accuracy than the less important regions (*ρ*_3*D–RC*_ < 0.1). Panels (C) and (D) illustrate dependency of regions with *E_local_*(***λ***)^-1^ ≥ 4 on iteration No. for pMLF1/14-3-3*ε* and npMLF1/14-3-3*ε*, respectively. Shown contours are *E_local_*(***λ***)^-1^ ≥ 4, which increases with proceeding iterations, and the contours covered the high-density regions (*ρ*_3*D–RC*_ ≥ 0.1) in the 14-th and 30-th iterations for pMLF1/14-3-3*ε* and npMLF1/14-3-3*ε*, respectively.

**Figure S7.**
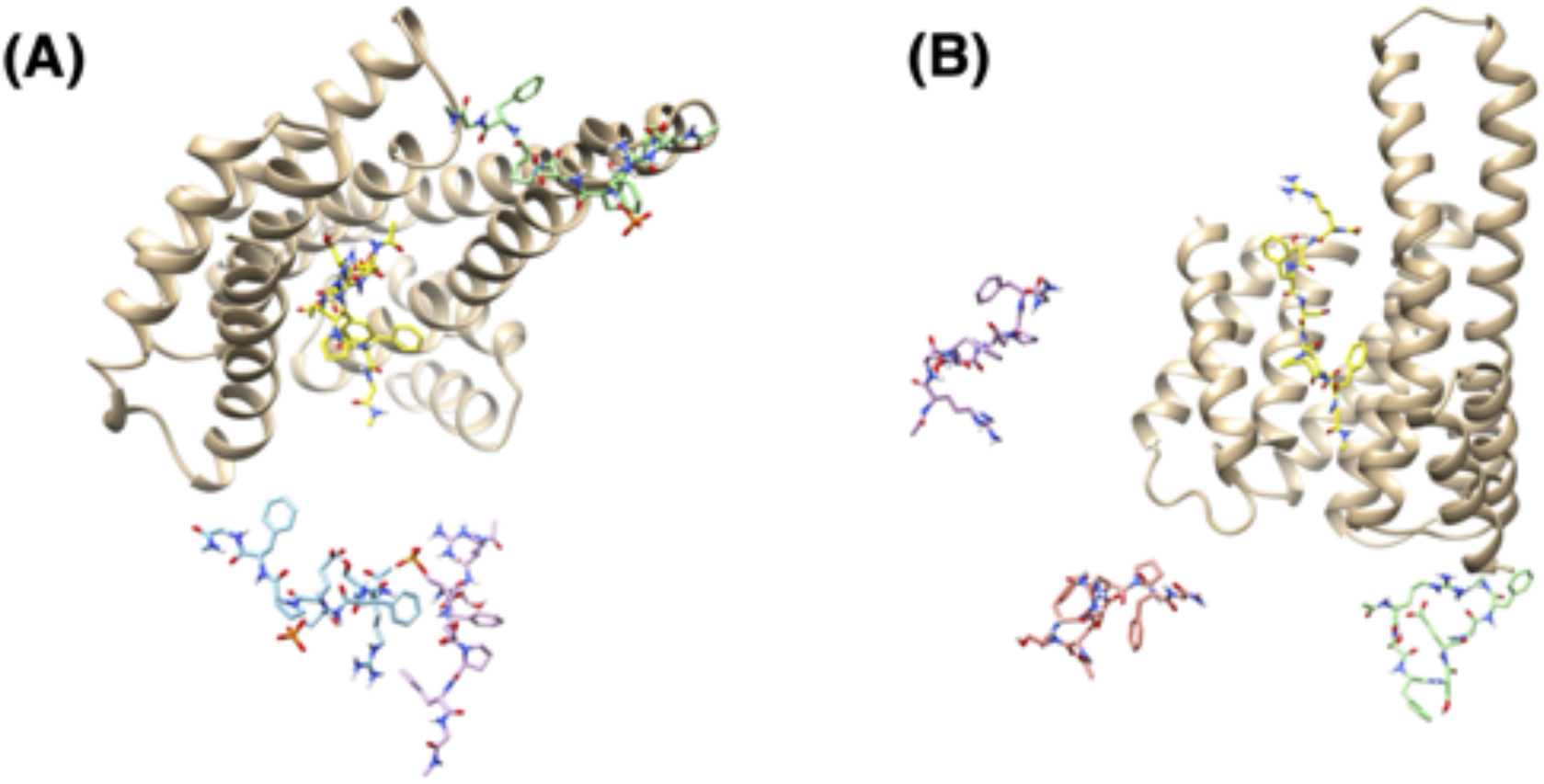
Panel (A) displays conformations taken randomly from cluster *C*_2_ of Fig. 2A for the pMLF1/14-3-3*ε* system, and panel (B) does those taken randomly from cluster 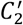 of Fig. 2B for the npMLF1/14-3-3*ε* system. Yellow-colored MLF1 conformations in the panels are those from crystal structure.

**Figure S8.**
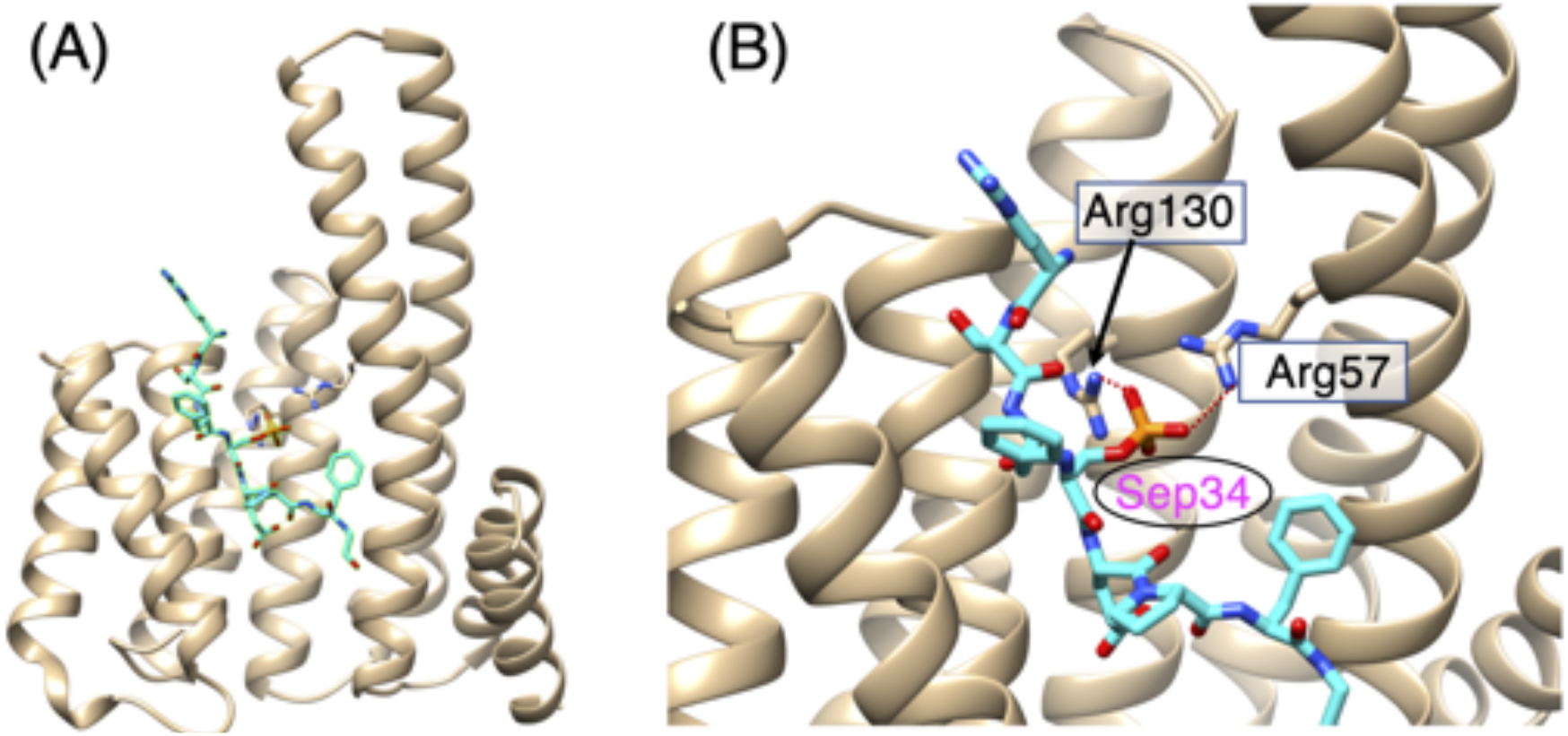
(A) pMLF1/14-3-3*ε* complex to show the position of Sep 34 in the entire complex structure. pMLF1 is shown by cyan-colored model, and 14-3-3*ε* by light yellow-colored model. (B) Close up of Sep 34 in pMLF1 and two residues Arg 57 and Arg 130 of 14-3-3*ε*, which form salt-bridges (brown-colored broken lines).

### Section 5. Calculation of radial distribution function

Here we present the method to calculate a radial distribution function. First, calculate a distance distribution function (or pair distance distribution function) *P*(*r^μ^*) (*μ* = *Phe*33, *Sep*34/*Ser*34, *Glu*35, or *Phe*37) from the equilibrated ensemble. A probability to detect the distance in a window [*r^μ^,r^μ^* + *dr^μ^*] is given by *P*(*r^μ^*)*dr^μ^*. See main text for the definition of distances *r^μ^*. Then, a radial distribution function (or radial density function) was defined as: *p*(*r^μ^*) = *P*(*r^μ^*)/4*π*(*r^μ^*)^2^ with normalization of ∫ *p*(*r^μ^*)*dr^μ^* = 1.

### Section 6. Calculation of spatial density of pMLF1/npMLF1 around 14-3-3*ε*

First, we divide the 3D real space into cubes, whose size Δ*V* is 2 Å × 2 Å ×2 Å. The cube position is defined by its center ***r*** = [*x, y, z*]. Then, we sum up the statistical weights of snapshots whose O*γ* atom of Sep 34 of pMLF1 (or Ser 34 of npMLF1) is detected in the cube. The sum of the weights is interpreted as a density at ***r***, and expressed as 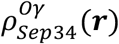 (or 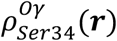). Last, we normalize the function as 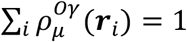, where the summation is taken over all cubes (∇*i*). This normalization makes the density map comparable between the pMLF1/14-3-3*ε* and npMLF1/14-3-3*ε* systems.

Other spatial density functions were calculated with the same manner. For instance, a density, 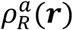, for the position of an atom *a* of residue *R* is calculated by summing up the statistical weights of snapshots whose atom *a* is detected in the cube ***r***. We use this spatial distribution function to analyze the distribution of Sep/Ser 34 and Glu 35 in this study. To analyze the density of Phe 33 and Phe 37, we calculated the position of the sidechain ring center of Phe 33 or Phe 37, and calculated the density functions for the ring centers: 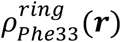 and 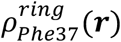.

**Figure S9.**
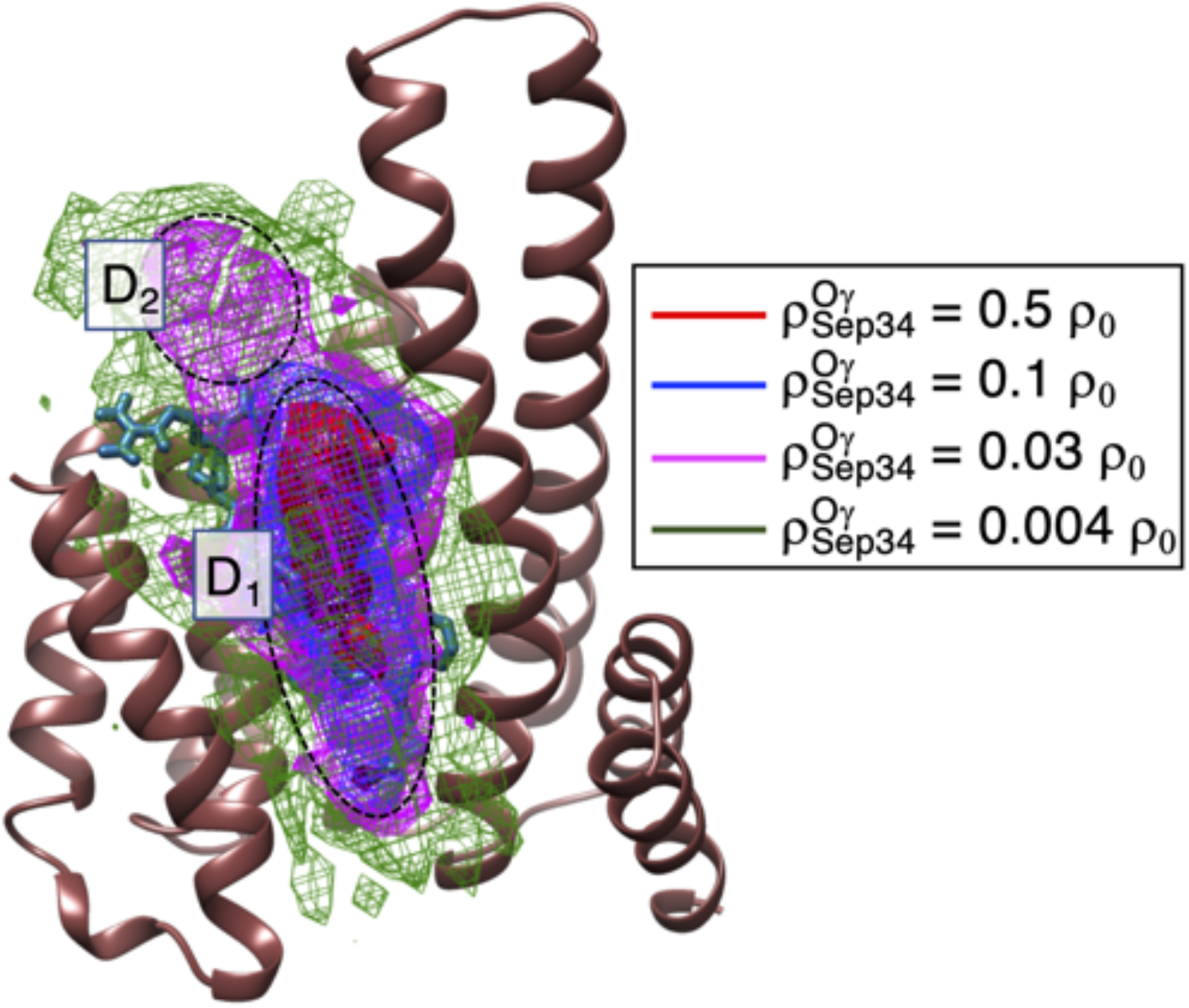
Distribution 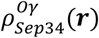 of the pMLF1/14-3-3*ε* system using snapshots in cluster *C*_1_, which was introduced in Fig. 2A of main text. Selection of snapshots from *C*_1_ was done as follows: Full variable ranges of RCs for this simulation are [6 Å,40 Å], [6 Å,40.5 Å], and [6 Å,40 Å] for *λ*_1_, *λ*_2_, and *λ*_3_, respectively (see Table S3 of SI). We verified that restricted ranges 6 Å≤ *λ*_1_ ≤ 20 Å, 6 Å≤ *λ*_2_ ≤ 20.25 Å, and 6 Å ≤ *λ*_1_ ≤ 20 Å do not involve snapshots in cluster *C*_2_. Thus, this figure was generated using the snapshots in the restricted ranges. This means that this figure was contributed by snapshots in intermediate regions other than *C*_1_ or *C*_2_. However, such conformations have a very small thermodynamics weights and do not contribute substantially to the shown distribution.

**Figure S10.**
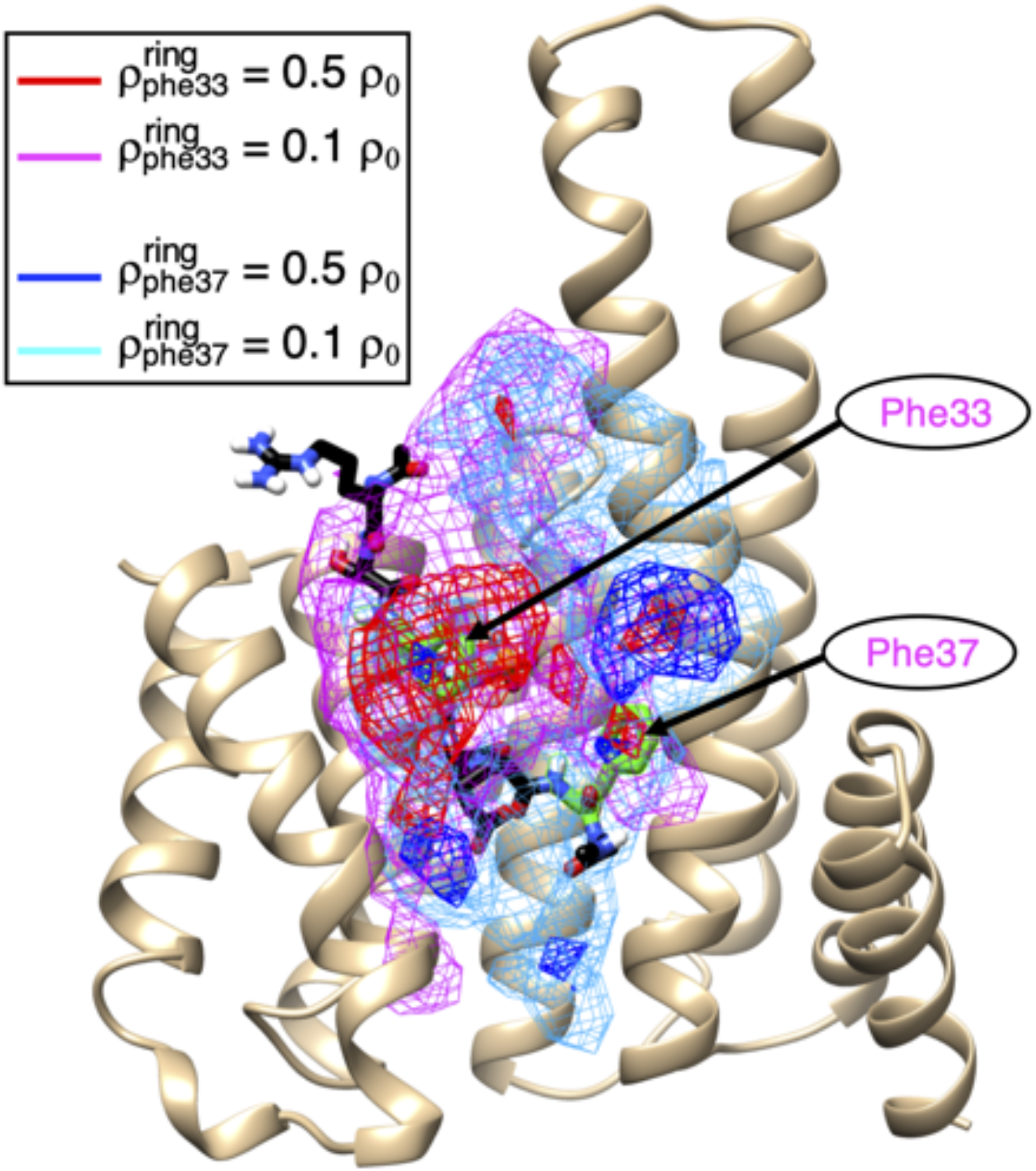
Distribution of 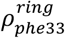 and 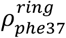 of the pMLF1/14-3-3ε system using snapshots only from cluster *C*_1_. See caption of Fig. S7 for selection of snapshots from *C*_1_.

**Figure S11.**
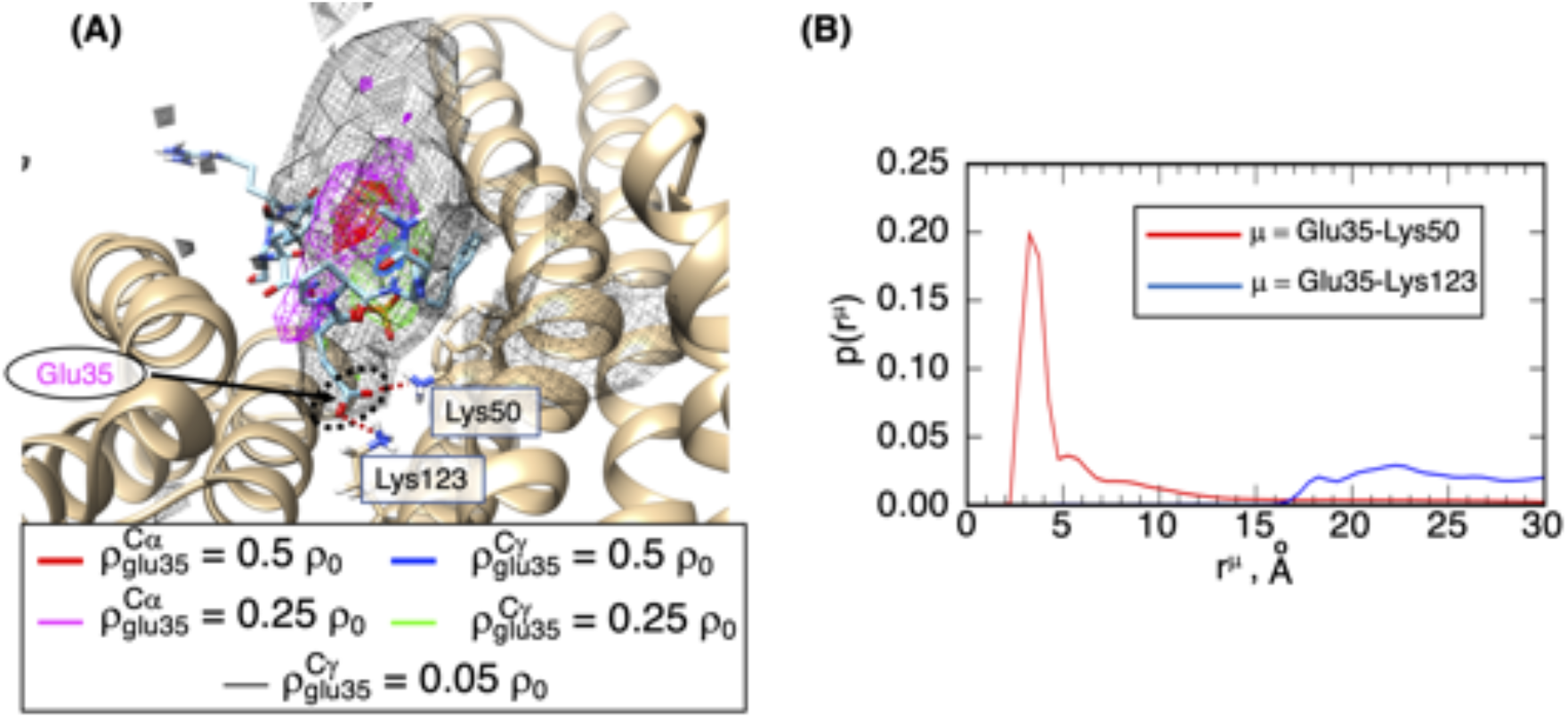
(A) Density contour maps of 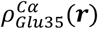 and 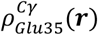 for pMLF1/14-3-3*ε*. Shown molecular structure is the crystal one focusing on Glu 35 of pMLF1. Contour levels are shown in inset (*ρ*_0_ = 0.01). Black-colored contours 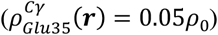 cover the sidechain-tip position of Glu 35 of the structure, where Glu 35 forms salt bridges (brown-colored broken line) to both Lys 50 and Lys 123 of 14-3-3*ε*. Black broken-line circle is mentioned in main text. (B) Radial distribution function, *p*(*r*^*Glu*35–*Lys*50^), for distance *r*^*Glu*35–*Lys*50^ from the Cγ atom of Glu 35 of pMLF1 to the N*ζ* atom of Lys 50 of 14-3-3 *ε*, and radial distribution function *p*(*r*^*Glu*35–*Lys*123^) for distance *r*^*Glu*35–*Lys*123^ from the C*γ* atom of Glu 35 to the N*ζ* atom of Lys 123 of 14-3-3*ε*. Method to calculate the radial distribution function is given in section 5 of SI.

### Section 7. Salt-bridge-formation probabilities

The salt-bridge-formation probabilities are defined by 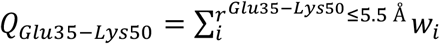 and 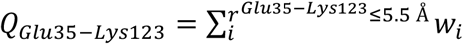, respectively. *r*^*Glu*35–*Lys*50^ is the distance from the C*γ* atom of Glu 35 of pMLF1 to the N*ζ* atom of Lys 50 of 14-3-3*ε*, and *r*^*Glu*35–*Lys*123^ is the distance from the C*γ* atom of Glu 35 to the N*ζ* atom of Lys 123 of 14-3-3*ε*. The *w_i_* is the statistical weight assigned to snapshot *i*. The summation for *Q*_*Glu*35–*Lys*50_ is taken over snapshots with *r*^*Glu*35–*Lys*50^ ≤ 5.5 Å and that for *Q*_*Glu*35–*Lys*123_ is done over those with *r*^*Glu*35–*Lys*123^ ≤ 5.5 Å. One may consider that the value of 5.5 Å is larger than a limit distance for a salt bridge. However, the C*γ* atom, not O*ε* atoms, was used to calculate *r*^*Glu*35–*Lys*50^ and *r*^*Glu*35–*Lys*123^, and Glu 35 has large positional fluctuations as shown above: Remember that a snapshot was saved once every 200 ps. Thus, we increased the limit distance.

### Section 8. Average of the orientation at each cube

In the main text, the orientation vector ***e**_i_* (|***e**_i_*| = 1) was defined for snapshot *i*. Here, we present a method for averaging the orientation vectors in each cube. As done for calculating the spatial density function (see section 6 of SI), we divided the 3D real space into cubes of 2.0 Å×2.0 Å×2.0 Å, whose positions are assigned to the cube centers ***r***. A snapshot was assigned to a cube when the O*γ* atom of Sep 34 of the snapshot is involved in the cube. Then, average of orientation vectors is taken for each cube:

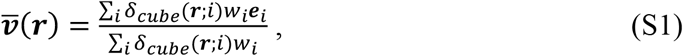

where *δ_cube_*(***r***; *i*) is a delta function defined as:

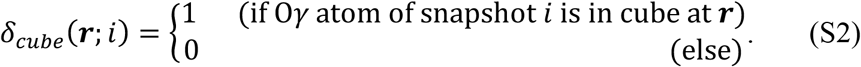

The quantity ∑_*i*_ *δ_cube_*(***r***; *i*)*w_i_* in Eq. S1 is equivalent to 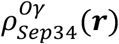. Note that 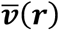 is no longer a unit vector. The norm of vector 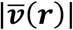 indicates a degree of orientational ordering of pMLF1 detected in cube 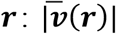 takes the maximum (= 1.0) when all ***e**_i_* have exactly the same orientations in the cube. If snapshots have uncorrelated orientations, then 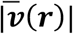 becomes small.

### Section 9. Ligand-conformational variety of pMLF1 at each cube

Here we present the procedure to calculate the ligand-conformational variety, Δ*r*_*Phe–Phe*_(***r***), of pMLF1 at each cube ***r***. First, we introduce a distance *r_Phe–Phe_*, which is the *Cα* atomic distance between Phe 33 and Phe 37. The value of *r_Phe–Phe_* for the crystal structure was 11.726 Å. Then we represent the ligand-conformational variety Δ*r_Phe–Phe_* by the standard deviation of *r_Phe–Phe_* in each cube ***r***:

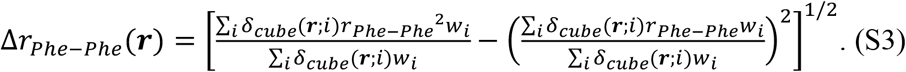

The larger the Δ*r*_*Phe–Phe*_(***r***) in a cube ***r***, the larger the ligand–conformational variety in the cube. We note that Δ*r*_*Phe–Phe*_(***r***) is not directly linked to a dynamic chain flexibility of pMLF1 because the conformational ensemble from the GA-guided mD-VcMD is an equilibrated one at 300 K.

**Figure S12.**
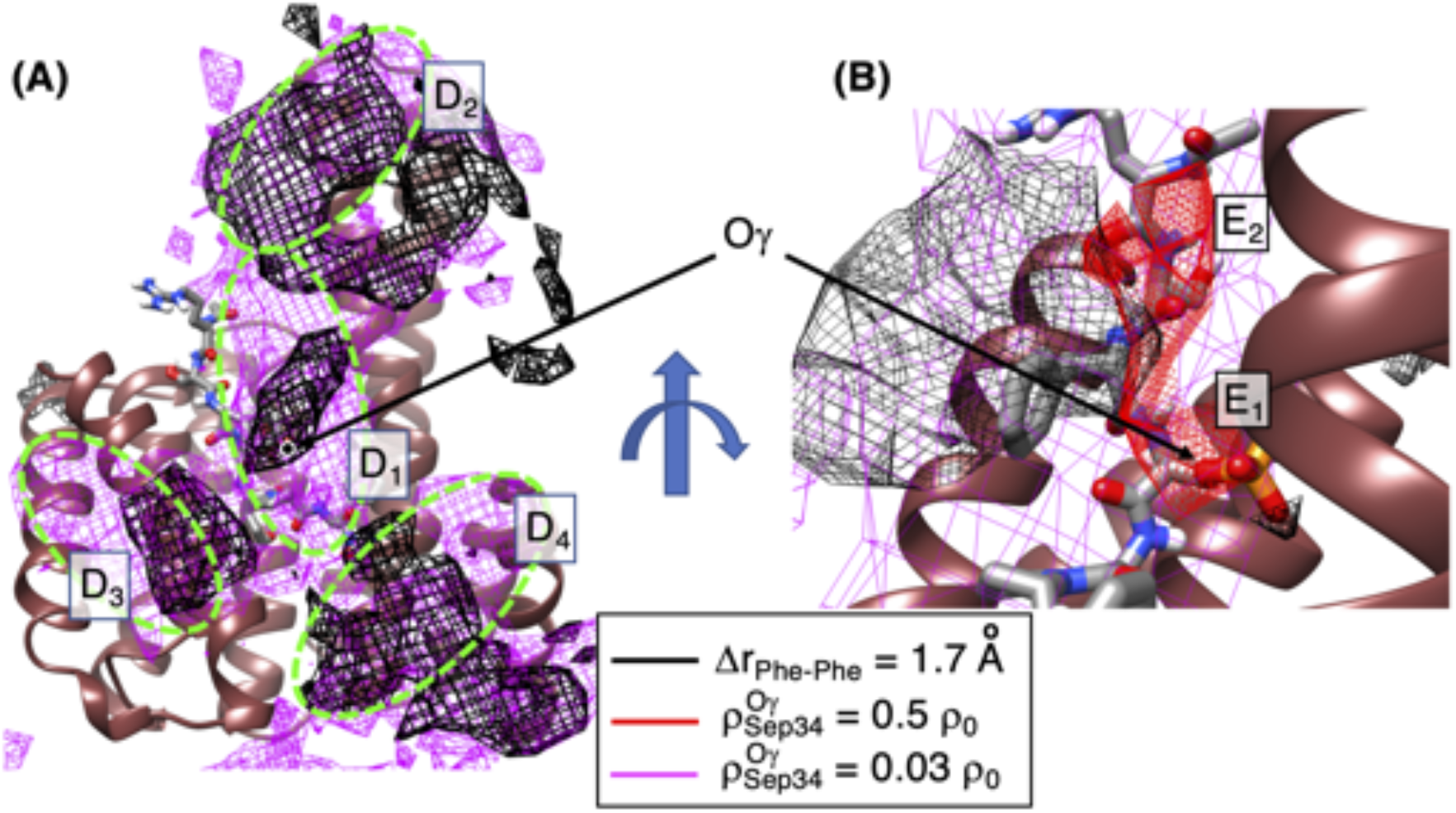
Density map 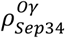 of pMLF1 and the large-Δ*r_Phe–Phe_* contours. Panel A, which is the same as Fig. 9B of main text except for the green-colored contours of Fig. 9B, shows clusters *D*_1_,…, *D*_4_. Panel B is a side view of panel A with focusing cluster *D*_1_. High-density spots *E*_1_ and *E*_2_ were defined in Fig. 6 of main text. Both panels present the position of *O_γ_* atom of Sep 34 in pMLF1.

### Section 10. Thermal average of distances in each cube

Here, we define a procedure to calculate thermal average of distances *r*^μ^ (*μ* = *Phe*33, *Sep* 34, *Glu*35, *Phe*37) by the following equation:

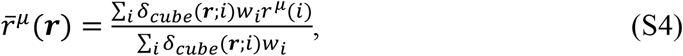

Where *r*^μ^(*i*) is *r^μ^* of snapshot *i*. The delta function *δ_cube_* (Eq. S2 in SI) works so that only snapshots, whose O*γ* atom of Sep 34 are detected in the cube at ***r***, contribute to 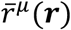. The 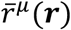 quantifies space dependency of the contacts of the four residues (Phe 33, Sep 34, Glu 35, and Phe 37) of pMLF1 to their contact partners of 14-3-3*ε* as a function of ***r***. The smaller the distance *r^μ^* in a cube ***r***, the contact tends to be formed in the cube.

